# TRAF2, an innate immune sensor, reciprocally regulates mitophagy and inflammation to maintain cardiac myocyte homeostasis

**DOI:** 10.1101/2021.06.12.448118

**Authors:** Xiucui Ma, David R. Rawnsley, Attila Kovacs, Moydul Islam, John T. Murphy, Chen Zhao, Minu Kumari, Layla Foroughi, Haiyan Liu, Krzysztof Hyrc, Sarah Evans, Brent A. French, Kenneth B. Margulies, Ali Javaheri, Babak Razani, Douglas L. Mann, Kartik Mani, Abhinav Diwan

## Abstract

Mitochondrial damage triggers cell death signaling with catastrophic consequences in long-lived and irreplaceable cells, such as cardiac myocytes. Sensing of leaked mitochondrial DNA upon mitochondrial damage is also a potent trigger of inflammation. Whether the innate immune response pathways monitor mitochondrial damage in mitochondria-rich cardiac myocytes to prevent inflammation and cell death, remains unknown. TRAF2, an adaptor protein downstream of innate immune receptors, localizes to the mitochondria in the unstressed heart, with increased mitochondrial targeting in cardiomyopathic human hearts and after cardiac ischemia-reperfusion injury in mice. Inducible cardiomyocyte-specific deletion of TRAF2 in young adult mice impairs mitophagy with rapid decline in mitochondrial quality, upregulates TLR9 expression in cardiac myocytes, and results in inflammation and cell death manifesting as a fulminant cardiomyopathy. Preventing TLR9-mediated mitochondrial DNA sensing and resultant inflammation provides a short-term reprieve from cardiomyopathy, but persistence of damaged mitochondria results in long-term recrudescence. Restoration of wild-type TRAF2, but not the E3 ubiquitin ligase deficient mutant, improves mitochondrial quality and rescues cardiomyopathy to restore homeostasis. Thus, the innate immune response acts via TRAF2 as the first line of defense against mitochondrial damage by orchestrating homeostatic mitophagy to dampen myocardial inflammation and prevent cell death.

## Introduction

Mitochondrial permeabilization results in activation of cell death signaling, and is a major pathway for cardiac myocyte death under stress.^1^ Damaged mitochondria also leak mitochondrial DNA, which is a potent trigger for immune activation due to its physical features that mimic bacterial DNA.^1^ Indeed, mitochondrial damage resulting in sterile myocardial inflammation and cardiac myocyte death is implicated in causing cardiomyopathy under pressure overload stress^2, 3^ and with ischemia-reperfusion injury^4, 5^; whereby targeting cell death as well as deleterious myocardial inflammation are regarded as key therapeutic targets to prevent and treat cardiomyopathy and heart failure.^1, 6^ Mitophagy is a lysosomal process to sequester damaged mitochondria (or parts thereof) to facilitate their removal via lysosomal degradation; and serves as a defense mechanism against mitochondrial DNA sensing to drive inflammation and cell death.^2, 7^ Whether the innate immune system responds to mitochondrial damage as a first responder to trigger mitophagy and suppress sterile inflammation in the myocardium, remains unknown.

Current understanding of the significance of mitophagy in the heart is predicated upon its role in removing damaged mitochondria and restoring normal mitochondrial quality primarily under stress, when mitochondria are damaged.^8–11^ While studies also point to a critical need for cardiac myocyte mitophagy during physiologic growth in the perinatal period^12^ or in the aging heart^13–15^; its role in cellular homeostasis in hearts of young mice is not well defined. Indeed, studies with novel and specific reporters have indicated robust rates of mitophagy in the unstressed adult mouse myocardium;^16, 17^ but ablation of previously characterized molecular mediators of mitophagy such as PINK1 and Parkin^18^ do not alter homeostatic mitophagy in the young mouse heart^8, 19–22^. In contrast, stressors such as ischemia-reperfusion injury,^8^ pressure overload hypertrophy^9, 23^ and metabolic stress with high fat diet feeding^10^ induce mitophagy; and Parkin ablation induces cardiac dysfunction indicating the mechanistic role of the PINK1-Parkin pathway in stress-induced mitophagy in cardiac myocytes. Accordingly, mechanisms and significance of homeostatic mitophagy in the young mouse heart remain to be defined.

Examination of mitochondrial turnover in cardiac myocytes reveals a half-life of ∼6-17 days as evaluated by conventional pulse chase assays^24, 25^, and contemporary analyses with maturation of mitoTimer fluorophore reveals that mitochondrial turnover slows down in the aging heart^26^. While mitochondrial protein quality control may proceed via multiple pathways,^27^ turnover of mitochondria requires the autophagy-lysosome pathway by mitophagy^18^. Indeed, impairment of global macro-autophagy with inducible adult-onset cardiac myocyte-specific ablation of ATG5^28^ or ATG7^29^, proteins essential for early steps in autophagosome formation, induces cardiomyopathy with appearance of damaged mitochondria on ultrastructural analyses. Taken together, these observations suggest that mitophagy is critical for mitochondrial quality control in the young adult heart in the unstressed state, a premise that requires experimental proof.

Prior studies also suggest that lysosome function is critical for preventing myocardial inflammation, as ablation of lysosomal DNase II, which degrades mitochondrial DNA, in cardiac myocytes, exacerbates myocardial inflammation to accelerate development of cardiomyopathy with pressure overload.^2^ Whether basal mitophagy suppresses myocardial inflammatory signaling in the unstressed heart remains unknown. It is noteworthy that all currently defined mitophagy pathways are intrinsic to damaged mitochondria and recognize mitochondrial damage or dysfunction to facilitate lysosomal removal of damaged mitochondria via mitophagy or other mechanisms;^27^ whereby participation of innate immune mechanisms in this process remains a missing piece of this conceptual framework. In this manuscript, we uncover evidence that TRAF2, an innate immunity effector that we have previously identified as a mediator of mitophagy in isolated cardiac myocytes,^30^ acts as a first line of defense against cardiac myocyte cell death and myocardial inflammation by facilitating mitophagy and suppressing TLR9 expression, respectively, *in vivo* in the mammalian heart. Our data demonstrate that a fraction of TRAF2 is associated with mitochondria in homeostasis; and its recruitment to the mitochondria is upregulated under conditions of mitochondrial damage with ischemia-reperfusion injury and in cardiomyopathic failing human hearts. Inducible ablation of TRAF2 in adult cardiac myocytes impairs mitophagy and provokes accumulation of damaged mitochondria, primarily based upon its role as an E3 ubiquitin ligase. Moreover, inducible TRAF2 ablation upregulates TLR9 expression to drive myocardial dysfunction and knockout of TLR9 delays the onset of cardiomyopathy by curtailing inflammation; but does not prevent eventual cardiomyopathic decompensation due to presence of damaged mitochondria. These studies highlight a central role for the innate immunity pathway as a mitochondria-extrinsic mechanism in sensing mitochondrial damage to effect mitophagy and maintain homeostasis.

## Results

### TRAF2 localizes to the mitochondria in human and mouse hearts

Our published studies in isolated primary neonatal rat cardiac myocytes indicate that TRAF2 localizes to mitochondria in an unstressed state, with increased localization upon injury.^30^ Moreover, prior work has demonstrated that TRAF2 localizes to the mitochondria in response to inflammatory stimuli in other cell types in mice.^31, 32^ To determine if this is also observed in human hearts, we obtained heart tissue from donors rejected for transplantation, as well as from patients with ischemic cardiomyopathy, and examined TRAF2 expression following biochemical fractionation into a mitochondria-enriched fraction and cytosol (Fig. 1a-d). TRAF2 expression was detected in the mitochondrial fraction (identified by enrichment for a mitochondrial protein, VDAC), where it was dramatically upregulated in hearts with ischemic cardiomyopathy and end-stage heart failure (Fig. 1b, d). This was accompanied by an increase in overall TRAF2 levels (Fig. 1e, f) as well as a relative decline in cytosolic levels (Fig. 1a, c), suggesting that cytosolic TRAF2 translocates to the mitochondria upon injury.

**Figure 1:**
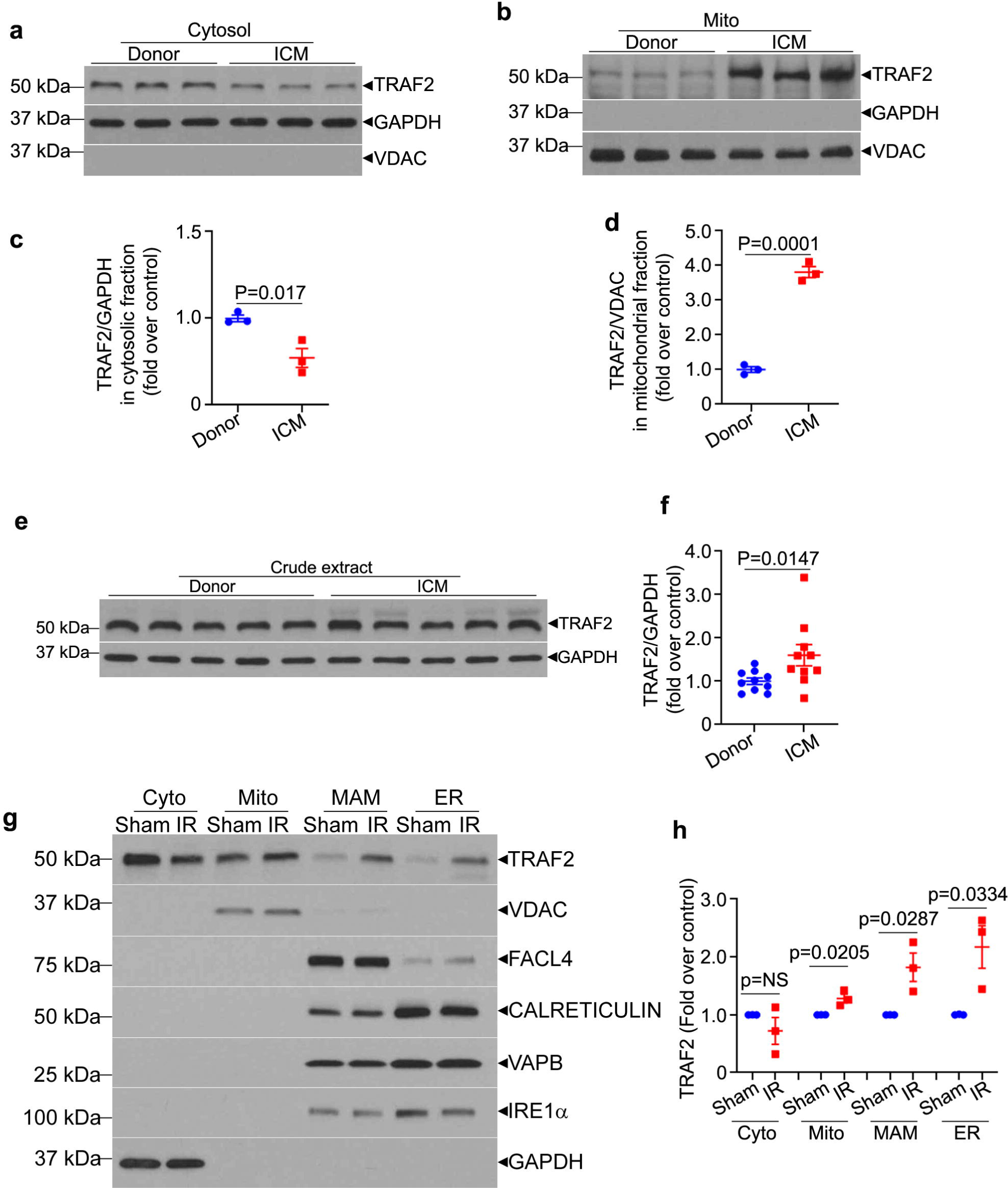
TRAF2 localizes to the mitochondria in human and mouse hearts. **a, b**. Immunoblots depicting TRAF2 expression in hearts from individuals evaluated as donors for cardiac transplantation, without evidence for cardiomyopathy (donor) or patients with end-stage ischemic cardiomyopathy (ICM) undergoing cardiac transplantation. The hearts were subjected to biochemical fractionation into mitochondria-enriched (b) and cytosolic fractions (a), shown by segregation of VDAC, a mitochondrial protein and GAPDH, a cytosolic protein, respectively. **c, d.** Quantitative assessment of TRAF2 expression in the respective biochemical fractions shown in a, b. **e, f**. Representative immunoblot and quantitation depicting total TRAF2 expression in crude extracts from humans hearts from patients with ischemic cardiomyopathy (ICM) or donors as control. **g, h**. Representative immunoblot depicting TRAF2 expression in hearts from male C57BL6J mice (8 weeks old) subjected to ex-vivo ischemia-reperfusion injury (IR) or sham (S) treatment as control. Expression of VDAC, FACL4, CALRETICULIN, VAPB and IRE1α is shown to evaluate co-segregation with mitochondria (mito, with VDAC), mitochondria-associated membranes (MAM, with FACL4), endoplasmic reticulum (ER, with CALRETICULIN, VAPB and IRE1α) and cytosol (cyto, with GAPDH). All P values shown are by t-test.

To critically examine the subcellular compartments that TRAF2 localizes to in the mouse heart, we subjected wild-type mouse hearts to ex-vivo ischemia-reperfusion injury (or sham treatment) and performed subcellular fractionation to further isolate the mitochondrial biochemical fraction into its components, namely pure mitochondria (without contamination from other organelles), mitochondria-associated membranes (MAM) and endoplasmic reticulum (ER).^33^ Our data (Fig. 1g, h) demonstrate that TRAF2 is predominantly located in pure mitochondrial fraction in the sham (unstressed) heart, with lesser degree of localization to the MAM and ER. Ex-vivo ischemia-reperfusion injury increased TRAF2 localization to all these compartments (Fig. 1g, h). Indeed, TRAF2 was also detected by immuno-gold analysis and transmission electron microscopy on mitochondrial membranes and MAM in HEK293 cells, with increased localization to these structures as well as to mitochondria in autophagosomes after hypoxia-reoxygenation injury (Fig. S1). These observations suggest that stress-induced increase in TRAF2 localization to mitochondria and associated structures may play a functional role in its cytoprotective effects, as suggested by prior studies from our group and others showing that TRAF2 is both necessary^34^ and sufficient^35–37^ to enhance cardiac myocyte survival after ischemia-reperfusion injury.

To examine the mechanisms for recruitment of TRAF2 to the mitochondria, we evaluated whether its mitochondrial localization depends upon PINK1. Indeed, as an executor of the canonical mitophagy pathway via macro-autophagy, PINK1, a serine-threonine kinase, is stabilized upon inner membrane depolarization following mitochondrial damage and recruits the E3 ligase Parkin.^38^ Our data demonstrate that PINK1 is not essential for recruitment of TRAF2 to the mitochondrial fraction under normoxic conditions, but markedly reduces TRAF2 translocation to the mitochondrial fraction upon hypoxia-reoxygenation injury (Fig. S2a, b, c) consistent with a lack of role for PINK1 in executing basal mitophagy in the unstressed heart.^19^ Taken together, these data indicate a role for TRAF2 in homeostatic mitochondrial quality control in cardiac myocytes, *in vivo*.

### TRAF2 is required for cardiac myocyte survival in the unstressed adult mouse heart

Prior studies have determined that perinatal cardiac myocyte-specific ablation of TRAF2 in the mouse heart provokes necroptosis of cardiac myocytes with systemic inflammation and cardiomyopathy.^34^ To isolate the effects of TRAF2 on cardiac myocyte homeostasis from developmental signaling, we examined the consequences of loss of TRAF2 in cardiac myocytes in the adult heart. Mice homozygous for floxed *Traf2* alleles (Traf2 fl/fl) carrying the *Myh6*-promoter driven Mer-Cre-Mer (MCM), were injected with tamoxifen (30 mg/kg/day) for 3 consecutive days to generate TRAF2-icKO (inducible cardiac-myocyte specific TRAF2 knockout) mice. Similarly treated TRAF2 fl/fl mice were studied as controls. TRAF2-icKO mice manifest symptoms of heart failure with reduced physical activity, hunched posture, labored breathing and ruffled coat within 5 days of initiating tamoxifen injections. Echocardiographic examination demonstrated marked left ventricular (LV) cavity dilation and reduced contractile performance (% fractional shortening (%FS); Fig. S3a). On necropsy, the hearts were enlarged with dilation of the LV cavity (Fig. S3b, c) and significantly increased heart weight (heart weight/tibial length ratio (mg/mm): 7.27±0.5 in TRAF2-icKO vs. 5.66±0.15 in Traf2 fl/fl, n=3/group; P=0.036). Immunoblot analysis demonstrated a 66% reduction in myocardial TRAF2 protein in TRAF2-icKO mice (Fig. S4a, b). While these data suggest that loss of TRAF2 results in cardiomyopathy, a decline in systolic performance was also detected in MCM transgenic mice as compared with wild-type alleles at the *Traf2* locus after tamoxifen injection (%FS: 30±4% vs. 51±4% in wild-type controls, P=0.02 by t-test. N=3/group).

As this signal of Cre-mediated toxicity as has been described previously,^39, 40^ we employed two other tamoxifen regimens that have been reported to circumvent Cre-mediated toxicity in the heart.^40, 41^ A 3 week regimen of (20mg/kg/day for 5 days per week) tamoxifen injections resulted in comparable knockdown of myocardial TRAF2 levels as compared with the 3-day regimen (Fig. S4c, d). As expected, isolation of adult cardiac myocytes separately from non-myocytes demonstrated near complete loss of TRAF2 protein in the cardiac myocytes (Fig. S4e, f) with this regimen in the TRAF2-icKO mice. Interestingly, TRAF2 protein expression was noted to be 3-fold higher in non-myocytes as compared with cardiac myocytes; and was not altered by MCM-mediated TRAF2 ablation (Fig. S4e, f). The mice were subjected to serial echocardiography at baseline, day 21 (after the last tamoxifen dose) and 2 weeks after the last dose was administered. We observed a strong trend towards increased mortality among TRAF2-icKO mice (5/13 vs. 0/10 in Traf2 fl/fl controls; P=0.065 by chi-square test) with signs of heart failure such as reduced activity, labored breathing and generalized swelling prior to death. None of the mice carrying the MCM transgene or Traf2 fl/fl mice demonstrated signs of heart failure or died during the study. We observed a decline in LV %FS (Fig. 2a) with LV dilation (Fig. 2b) in TRAF2-icKO mice as compared with Traf2 fl/fl controls, that persisted at follow up. We also observed a transient decline in %FS and mild LV dilation in MCM transgenic mice, which was completely reversed at day 35 (Fig. 2a, b), demonstrating evidence for transient Cre-mediated toxicity with this protocol. Surviving TRAF2-icKO mice demonstrated increased fibrosis at day 35 (Fig. 2c, d) and ultrastructural examination revealed abnormal swollen mitochondria with cristal rarefication (white arrows; Fig. 2e). This was accompanied by a ∼4-fold increase in TUNEL positive cardiac myocytes as compared with Traf2 fl/fl controls (Fig. 2f). MCM mice did not display fibrosis, TUNEL positivity, or ultrastructural abnormalities indicating the Cre toxicity was transient (Fig. 2c-f). Importantly, the myocardium in MCM transgenic mice was histologically indistinguishable from wild-type mice without increase in TUNEL positive cardiomyocytes (Fig. 2c-f). Taken together, these data indicate that the loss of TRAF2 in cardiac myocytes provokes cardiomyocyte death in unstressed hearts with spontaneous development of cardiomyopathy. To address the possibility that Cre-mediated toxicity may still be contributing to the observed cardiomyopathy in TRAF2-icKO, we employed another regimen of a single dose of tamoxifen (40mg/kg) followed by echocardiography 2 weeks later.^40^ This regimen induced only a modest TRAF2 protein knockdown in the myocardium (Fig. S4g, h) and resulted in a mild cardiomyopathy with mildly dilated LV (in mm: 3.6±0.1 in TRAF2-icKO vs. 3.3±0.1 in Traf2 fl/fl, P=0.017, N=4/group) with mild systolic dysfunction (%FS: 41±1 in TRAF2-icKO vs. 50±1 in Traf2 fl/fl, P<0.001). At this dose, we did not observe any cardiac structural or functional abnormalities in MCM transgenic mice as compared with wild-type controls.

**Figure 2:**
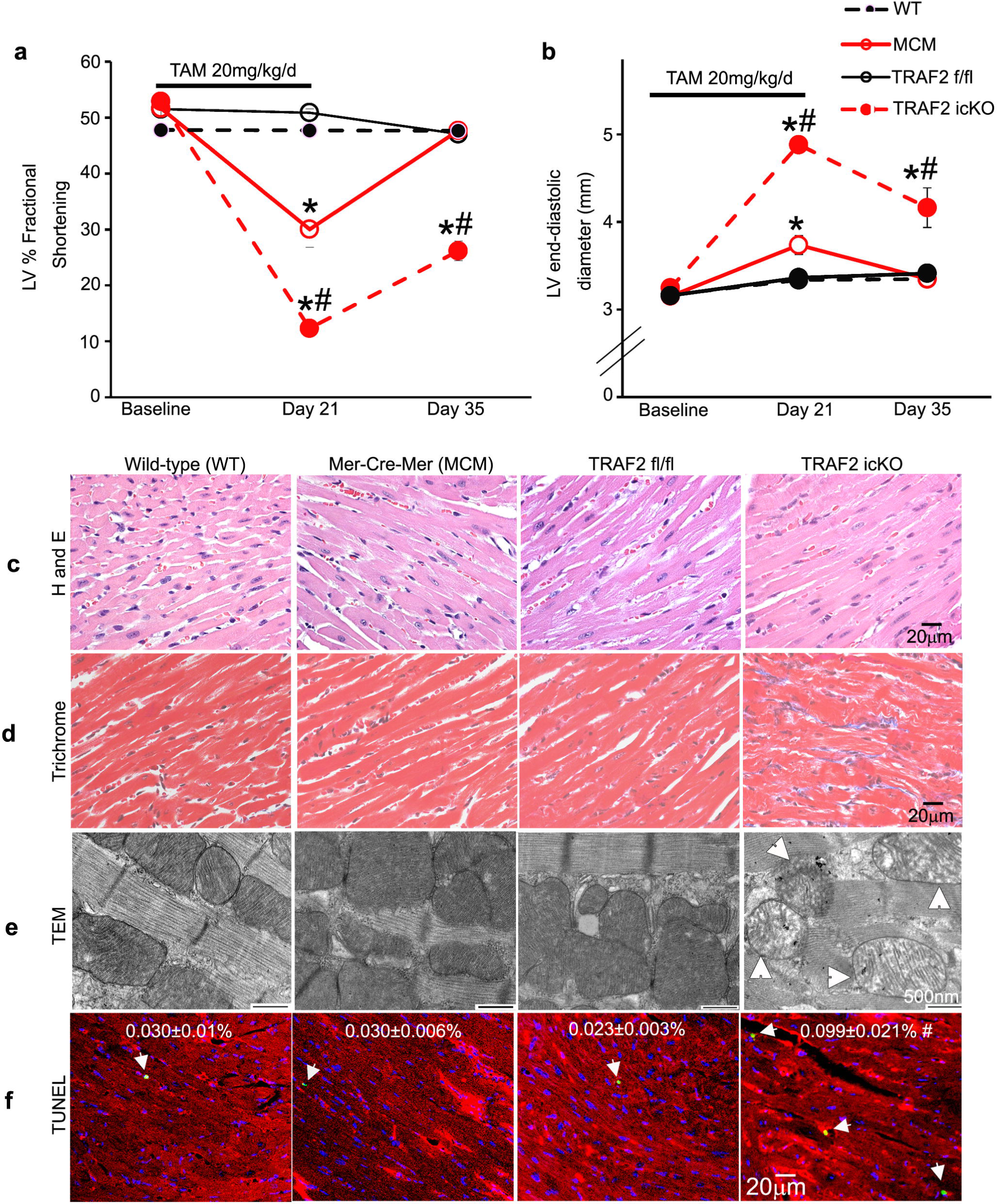
TRAF2 ablation in adult cardiac myocytes induces cardiomyopathy with damaged mitochondria and cell death. **a, b**. Left ventricular (LV) endocardial fractional shortening (%FS, a) and end-diastolic diameter (b) in wild-type, *TRAF2* floxed (Traf2 fl/fl), MerCreMer (MCM) mice, and mice homozygous for *TRAF2* floxed alleles carrying the MerCreMer transgene (Traf2 fl/fl+MCM), 14 days after tamoxifen treatment (20 mg/kg/d, i.p. 5 days per week for 3 weeks). N=9-10 mice/group. ‘*’ indicates P<0.05 versus wild-type and ‘#’ indicates P<0.05 versus Traf2 fl/fl by post-hoc test after two-way ANOVA. **c, d**. Representative images with hematoxylin and eosin staining (c), and trichrome staining (d) to evaluate myocardial structure and fibrosis, respectively, in mice as treated in a, b. **e.** Representative transmission electron microscopy images to evaluate myocardial ultrastructure in mice as in a. White arrows indicated swollen mitochondria with cristal rarefication. **f.** Representative TUNEL stained images to evaluate cardiac myocyte TUNEL positivity (green) in mice as in a. DAPI staining identifies nuclei. Staining for α-sarcomeric actin was performed to evaluate TUNEL staining in cardiac myocyte nuclei. %TUNEL positive nuclei/total cardiac myocyte nuclei are depicted. N=4-6/group. ‘#’ indicates P<0.05 versus Traf2 fl/fl by post-hoc test after one-way ANOVA.

Overall, we observed a strong correlation between the degree of TRAF2 protein knockdown in the myocardium in the TRAF2-icKO mice with various tamoxifen regimens; and the decline in % fractional shortening and the observed LV dilation (Fig. S4i, j). We also set up a model of inducible cardiac myocyte TRAF2 ablation by feeding tamoxifen chow. Mice treated with tamoxifen chow for 7 days demonstrated a similar degree of TRAF2 deletion as the 21-day injection regimen and with comparable LV dilation and systolic dysfunction as compared with floxed controls (Fig. S5a, b). These data demonstrate that inducible loss of TRAF2 in adult cardiac myocytes provokes cardiomyopathy with increased cardiac myocyte death. Prior studies indicate that perinatal onset ablation of TRAF2 in cardiac myocytes (with Myh6-Cre) induces cardiac myocyte necrosis with systemic inflammation.^34^ Accordingly, we assessed for circulating markers noted to be elevated with TRAF2 ablation in the prior work. Our data demonstrate that circulating HMGB1 levels, a marker of necrotic cell death (Fig. S6a) or circulating cytokines (IL1β, TNF and IL-6; Fig. S6b-d) are not elevated in TRAF2-icKO mice as compared with TRAF2 fl/fl controls; pointing to a different mechanism for the observed cardiomyopathy in mice with adult onset TRAF2 ablation.

### Ablation of TRAF2 impairs cardiac myocyte mitophagy in the adult mouse heart

We observed that mitochondrial proteins accumulate upon TRAF2 ablation in the TRAF2-icKO mice (Fig. 3a, Fig. S5b, d, e, f), pointing to impaired removal of damaged mitochondria with TRAF2 deficiency (as seen with TEM analysis in Fig. 2e). Indeed, examination of homeostatic mitophagy with AAV9-mediated transduction of mKeima reporter^16^ revealed marked suppression of cardiac myocyte autophagy in the TRAF2-icKO mice (Fig. 3b-d). Interestingly, levels of Parkin, a mitophagy mediator under stress, remained unchanged with inducible TRAF2 ablation (Fig. 3a). We next examined if the mitophagy deficit was secondary to defects in macro-autophagy, by examining flux in TRAF2-icKO mice expressing a dual fluorescent flux reporter. Our data demonstrate that the prevalence of autolysosomes relative to total autophagic structures, as well as abundance of autolysosomes and autophagosomes was not reduced in TRAF2-icKO as compared with TRAF2 fl/fl mice (Fig. 3e-g). These data indicate that the observed defect with TRAF2 ablation is with organelle-specific autophagy, i.e. mitophagy without impairment of macro-autophagy in TRAF2-icKO hearts.

**Figure 3:**
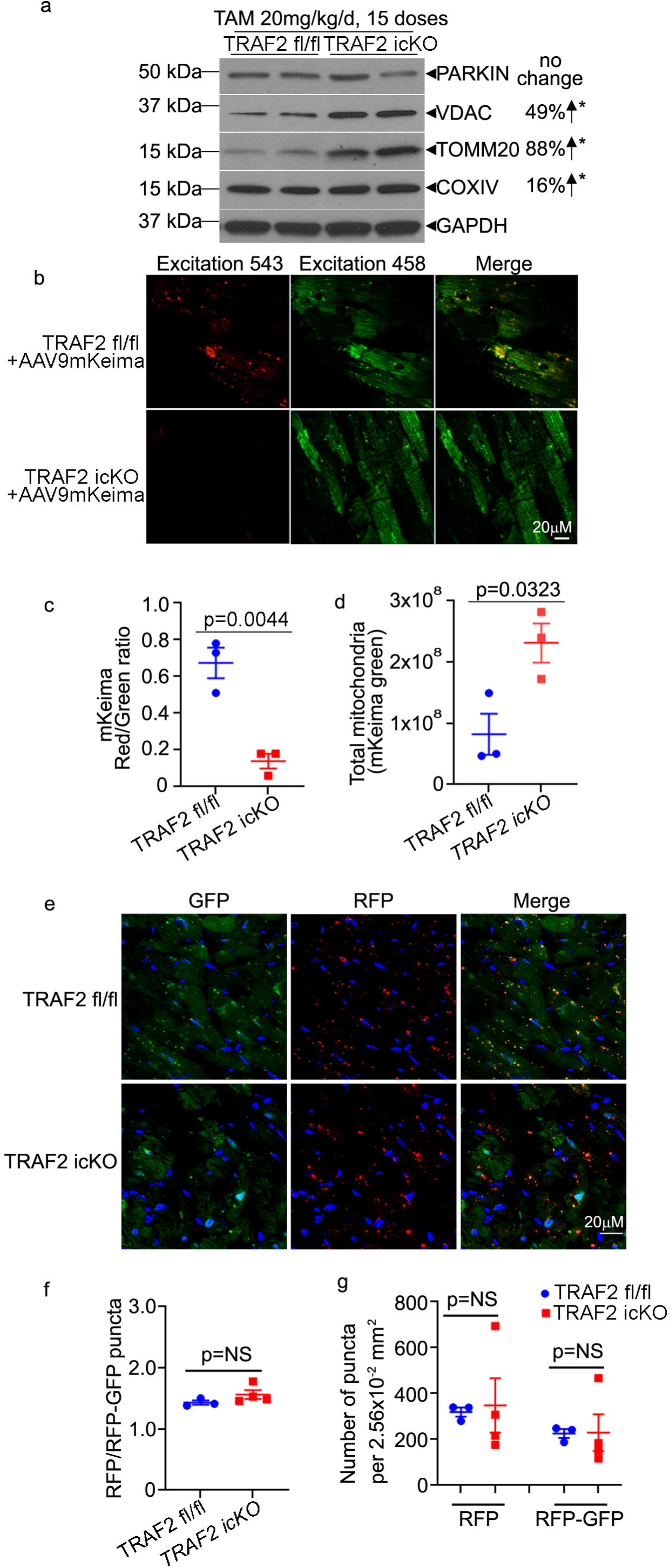
TRAF2 ablation in adult cardiac myocytes impairs homeostatic mitophagy. **a.** Representative immunoblot depicting expression of PARKIN and mitochondrial proteins (VDAC, TOMM20, COXIV) in cardiac extracts from mice at day 35 after inducible adult-onset ablation of TRAF2 (TRAF2-icKO), 14 days after tamoxifen treatment (20 mg/kg/d, i.p. for 5 days per week for 3 weeks). N=4/group. Protein levels were normalized to GAPDH. ‘*’ indicates P<0.05 for TRAF2-icKO versus TRAF2 fl/fl groups by t-test; and upward arrow points to upregulation in abundance of indicated protein indicated as a % change. **b-d**. Representative images (b) from hearts transduced with AAV9-mitoKeima in mice as in a; with ratio of quantitation of mitoKeima emission in the red channel over green as index of mitophagy (c). Total mitochondria were assessed by abundance of green signal normalized to the area of transduced cardiac myocytes. ‘*’ indicates P<0.05 by t-test for TRAF2-icKO versus TRAF2 fl/fl groups. **e-g.** Representative images (e) for assessment of flux through macro-autophagy in TRAF2-icKO versus TRAF2 fl/fl mice carrying the RFP-GFP-LC3 reporter transgene, with quantitative assessment of ratio of autolysosomes (punctate RFP signal) to autophagosomes (punctate RFP+GFP signal, in f); and total autophagosomes (RFP+GFP puncta) and autolysosomes (RFP puncta) per unit myocardial area in g. NS indicates P>0.05 by t-test.

### TRAF2 ablation results in mitochondrial DNA release and induction of TLR9 expression

Impairment of mitophagy results in mitochondrial DNA leak from damaged mitochondria, and prior studies have demonstrated TLR9-mediated mitochondrial DNA sensing is a driver of myocardial pathology under pressure overload stress.^2, 4^ Indeed, TRAF2-icKO myocardium demonstrated increased co-localization of DNA detected by PicoGreen with TLR9 in cardiac myocytes (Fig. 4a). Furthermore, TRAF2 ablation resulted in transcriptional upregulation of TLR9 expression (Fig. 4b-d), which was localized to cardiac myocytes (Fig. 4e) per in-situ hybridization to detect TLR9 transcript expression. These data indicate that loss of TRAF2 sensitizes the myocardium to pro-inflammatory signaling driven by TLR9 sensing of leaked mitochondrial DNA.

**Figure 4:**
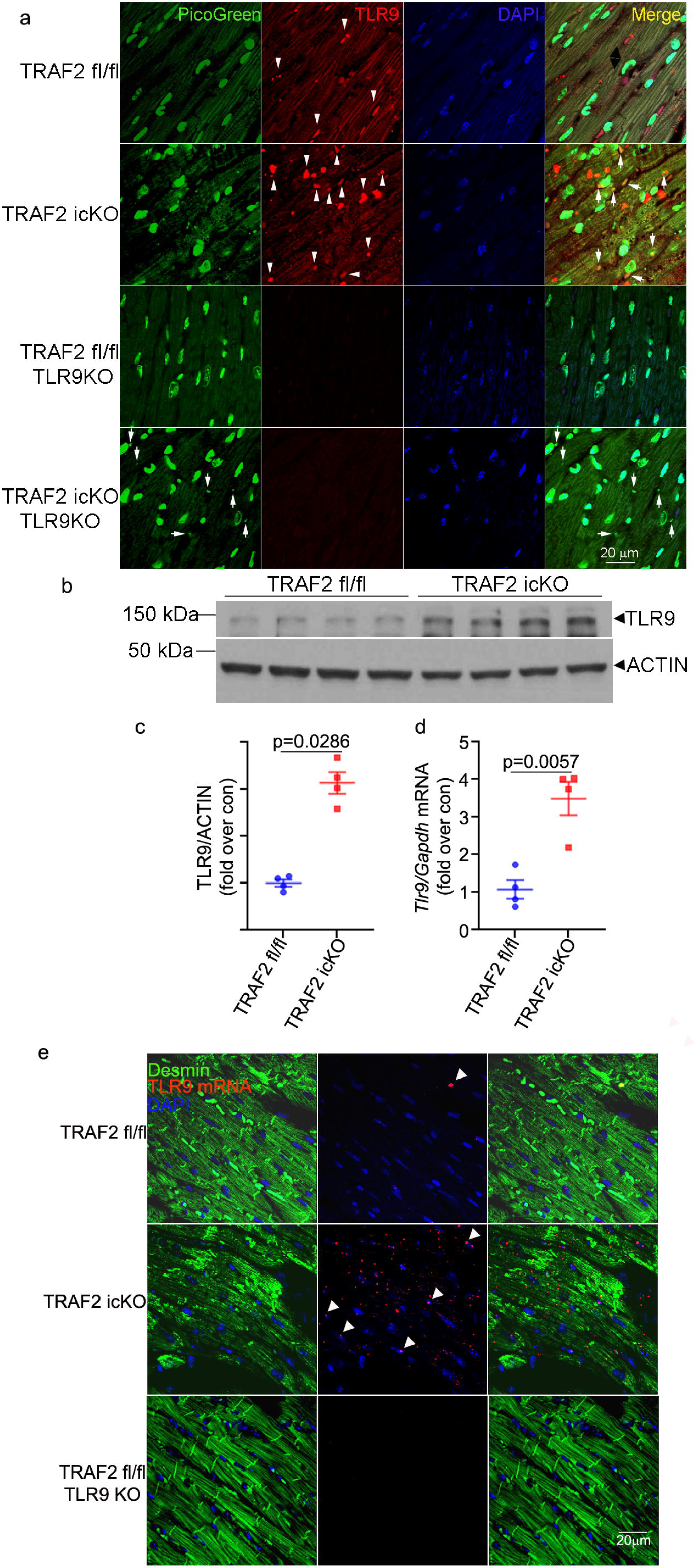
TRAF2 ablation in adult cardiac myocytes upregulates TLR9 and mitochondrial DNA sensing. **a.** Representative images demonstrating co-localization of PicoGreen with TLR9 in myocardial sections from mice with adult-onset inducible TRAF2 ablation (TRAF2-icKO) and TRAF2 floxed controls, without and with concomitant germline TLR9 ablation. Nuclei are stained blue (DAPI). White arrowheads demonstrate co-localization of PicoGreen with TLR9. White arrows point to cytosolic detection of DNA by PicoGreen in TRAF2-icKO-TLR9KO myocardium. **b, c**. Immunoblot (b) and quantitation (c) depicting TLR9 expression in cardiac extracts from TRAF2-icKO and TRAF2 floxed control mice modeled as in a. P value depicted is by t-test. **d**. *Tlr9* transcript expression in cardiac extracts from TRAF2-icKO and TRAF2 floxed control mice as in a. P value depicted is by t-test. **e**. In-situ hybridization to detect *Tlr9* transcript (red) and its co-localization with DESMIN (green) to identify cardiac myocytes. Arrowheads point to TLR9 transcripts in cardiac myocytes. Representative section from *Traf2* fl/fl *Tlr9* null mouse is shown as control.

### Concomitant TLR9 ablation rescues cardiomyopathy in the setting of cardiac myocyte TRAF2 loss

Mitochondrial damage triggers inflammation as well as cell death signaling. To determine whether inflammation observed in TRAF2-icKO myocardium is a direct cause of cardiac myocyte death, as described in prior studies that examined the effects of perinatal cardiac myocyte TRAF2 ablation^34^, we examined the prevalence of CD68+ macrophages. Our data demonstrate an increased in abundance of cardiac CD68+ cells (Fig. 5a, b) in TRAF2-icKO myocardium indicating cellular inflammatory infiltrate. Notably, concomitant ablation of TLR9 (Fig. S7a, b) markedly attenuated the abundance of cardiac CD68+ cells to levels observed in controls (Fig. 5a, b), as well in TUNEL+ cardiac myocytes in mice with inducible cardiac myocyte TRAF2 ablation (Fig. 5c, d) with reduced fibrosis and unaffected myocyte registration (Fig. S7c, d). Concomitant TLR9 ablation also attenuated the decline in systolic function (Fig. 5e) and left ventricular hypertrophy (Fig. 5g, h) without significantly affecting left ventricular dimensions (Fig. 5f) in TRAF2-icKO mice. This was accompanied by attenuated fetal gene expression pattern as a marker for pathologic hypertrophy (with reductions in *Anp*, *Bnp*, *Acta1* and restoration of *Serca2a* transcripts (Fig. 5i-l) without an effect on *Myh6* and *Myh7* expression (Fig. S7e, f) and a marked reduction in TUNEL positive cardiac myocytes (Fig. 5c, d). Taken together, these data suggest that TLR9-mediated inflammatory signaling in the myocardium drives cell death and myocardial pathology in TRAF2-icKO mice without a contribution from systemic inflammation driven by necroptotic cell death, as was observed with perinatal TRAF2 ablation.^34^

**Figure 5.**
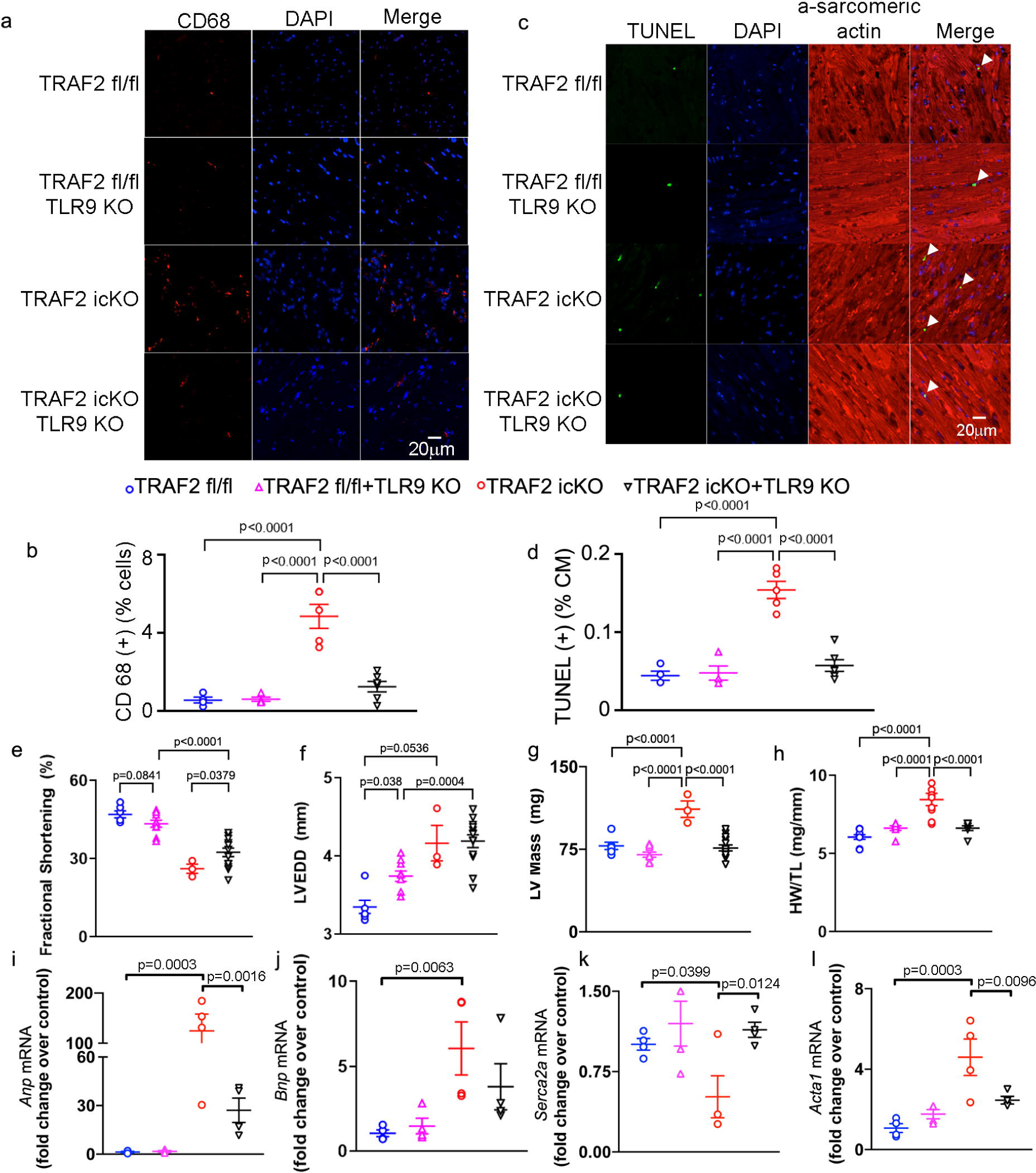
TLR9 ablation rescues inflammatory cell infiltration and cell death in mice with inducible TRAF2 ablation in adult cardiac myocytes, in the short term. **a.** Representative images depicting CD68+ cells in myocardial section from mice with adult-onset inducible TRAF2 ablation (TRAF2-icKO) and TRAF2 floxed controls, without and with concomitant germline TLR9 ablation, 2 weeks after TRAF2 ablation. Nuclei are stained blue (DAPI). **b**. Quantitation of CD68+ cells as % of all DAPI stained nuclei in mice as in a. P values by post-hoc test after one-way ANOVA. **c, d**. Representative TUNEL stained images (c) with quantitative assessment (d) of cardiac myocyte TUNEL positivity (green; %TUNEL positive nuclei/total cardiac myocyte (CM) nuclei) in mice as in a. DAPI staining identifies nuclei. Staining for α-sarcomeric actin was performed to evaluate TUNEL staining in cardiac myocyte nuclei. P values by post-hoc test after one-way ANOVA. **e-h**. Left ventricular endocardial fractional shortening (%FS, e), end-diastolic diameter (LVEDD, mm, f), mass (mg, g) and heart weight normalized to tibial length (mg/mm, h) in mice modeled as in a. P values are by post-hoc test after one-way ANOVA. **i-l**. Expression of *Anp* (i), *Bnp* (j), *Serca2a* (k), and *Acta1* (l) transcripts in mice modeled as in a. P values by post-hoc test after one-way ANOVA.

Evaluation of mitochondrial ultrastructure revealed that concomitant ablation of TLR9 in TRAF2-icKO mice did not rescue the mitochondrial defects observed with TRAF2 ablation in cardiac myocytes (Fig. 6a; also see Fig. 2e). Moreover, mitochondrial DNA leak in the cytosol was detectable by PicoGreen signaling in TRAF2-icKO mice with TLR9 ablation (Fig. 4a). This suggested that TLR9 activation is downstream of effects of TRAF2 ablation on the mitochondria. Damaged mitochondria trigger cell death signaling (reviewed in ^1^), which prompted us to follow these animals over a longer time-period to determine if the rescue of cardiac myocyte death by abrogation of TLR9-mediated DNA sensing was durable. Indeed, re-examination of these animals at 36 weeks after tamoxifen treatment (to induce TRAF2 ablation) resulted in LV dilation (Fig. 6b), LV systolic dysfunction (Fig. 6c), increased LV mass (Fig. 6d) and with increased ratio of cavity size to wall thickness (r/h, Fig. 6e) in TRAF2-icKO-TLR9 null mice; indicating that persistence of damaged mitochondria is sufficient to induce cardiomyopathy independent of sustained inflammatory signaling.

**Figure 6.**
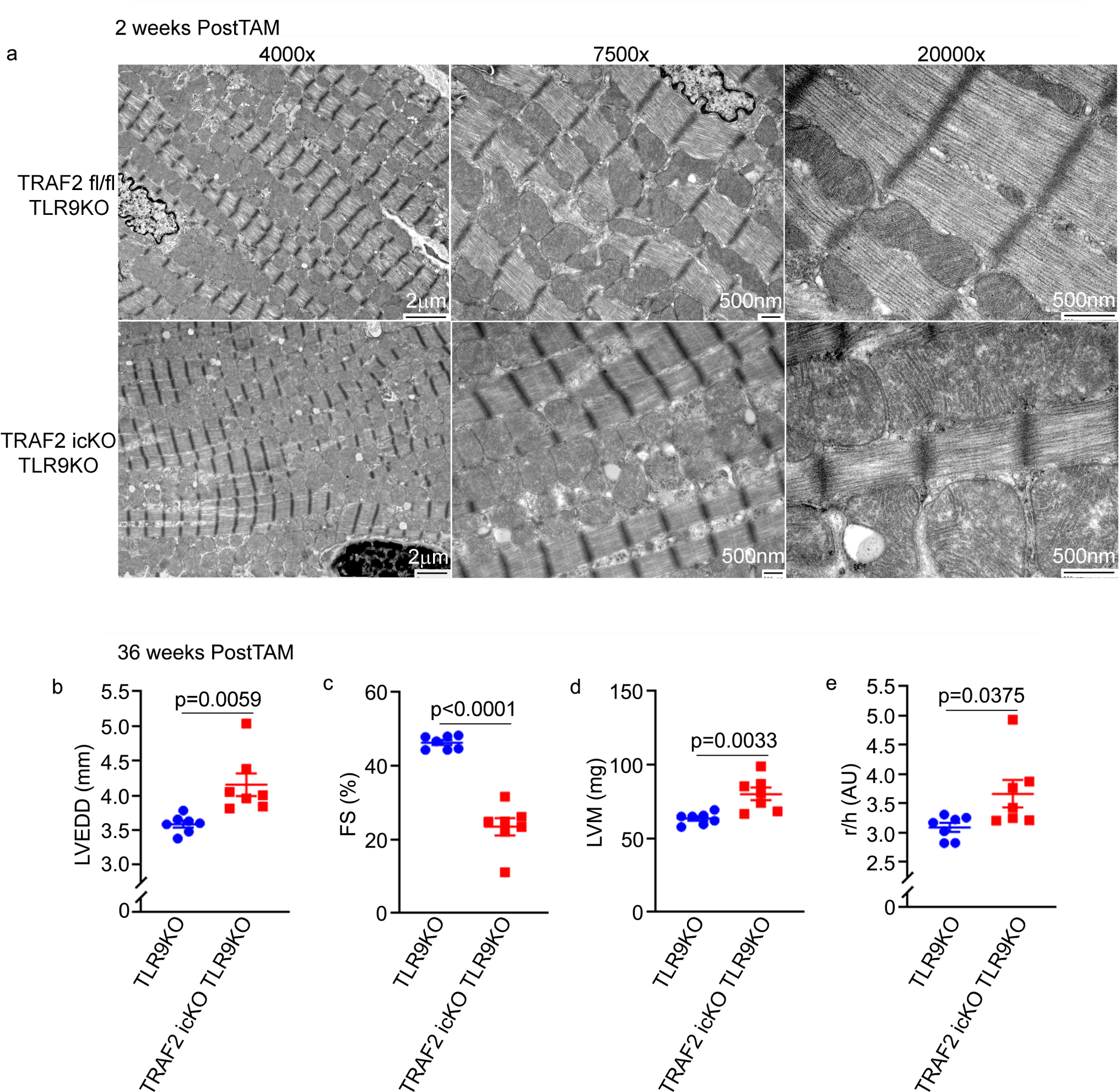
TLR9 ablation does not rescue mitochondrial abnormalities and does not prevent cardiomyopathy with inducible cardiac myocyte TRAF2 ablation with longer-term follow up. **a.** Representative transmission electron microscopy images to evaluate myocardial ultrastructure in mice as with adult-onset inducible TRAF2 ablation (TRAF2-icKO) and TRAF2 floxed controls, without and with concomitant germline TLR9 ablation, 2 weeks after TRAF2 ablation. **b-e**. Mice with adult-onset inducible TRAF2 ablation or TRAF2 floxed alleles in TLR9 null background (TRAF2-icKO TLR9-/- and TRAF2 fl/fl TLR9-/-) were followed by echocardiographic evaluation at 36 weeks after tamoxifen treatment to induce TRAF2 ablation. Left ventricular end-diastolic diameter (LVEDD, mm, b), endocardial fractional shortening (%FS, c), mass (mg, d) and ratio chamber diameter to wall thickness (r/h, e) was assessed as shown. P values shown are by t-test.

### AAV9-mediated restoration of TRAF2 but not its E3 ligase deficient mutant rescues cardiomyopathy with cardiac myocyte TRAF2 ablation

To examine the specificity of the observations with inducible adult-onset cardiac-myocyte ablation of TRAF2, and gain a mechanistic understanding of whether its E3 ligase function was critical as we have observed previously in cardiac myocytes in cell culture,^30^ we utilized a AAV9-based approach to restore TRAF2 expression in TRAF2-icKO myocardium. Indeed, AAV9-mediated expression of TRAF2 targeted to cardiac myocytes with a cTnT promoter^42^ rescued the decline in systolic function (Fig. 7a, b) and restored mitochondrial protein levels to those observed in mice without TRAF2 ablation (Fig. 7c, d). Notably, the mitochondrial morphology was also normalized in TRAF2-icKO mice with AAV9-mediated TRAF2 delivery (Fig. 7h). In contrast, AAV9-mediated restoration of TRAF2 with mutations in ring domain residues to ablate its E3 ligase activity (i.e. TRAF2-Rm mutant, as previously described^30, 43^; see Fig. 7c) did not rescue the systolic dysfunction (Fig. 7a, b), restore abundance of mitochondrial proteins (Fig. 7c, e-g) or rescue the mitochondrial defects (Fig. 7h). Taken together, these data demonstrating that the E3 ligase domain of TRAF2 is critical to its role in myocardial homeostasis and mitochondrial quality control, mirroring the observations with its critical role in mediating mitophagy in isolated primary cardiac myocytes in cell culture.^30^

**Figure 7.**
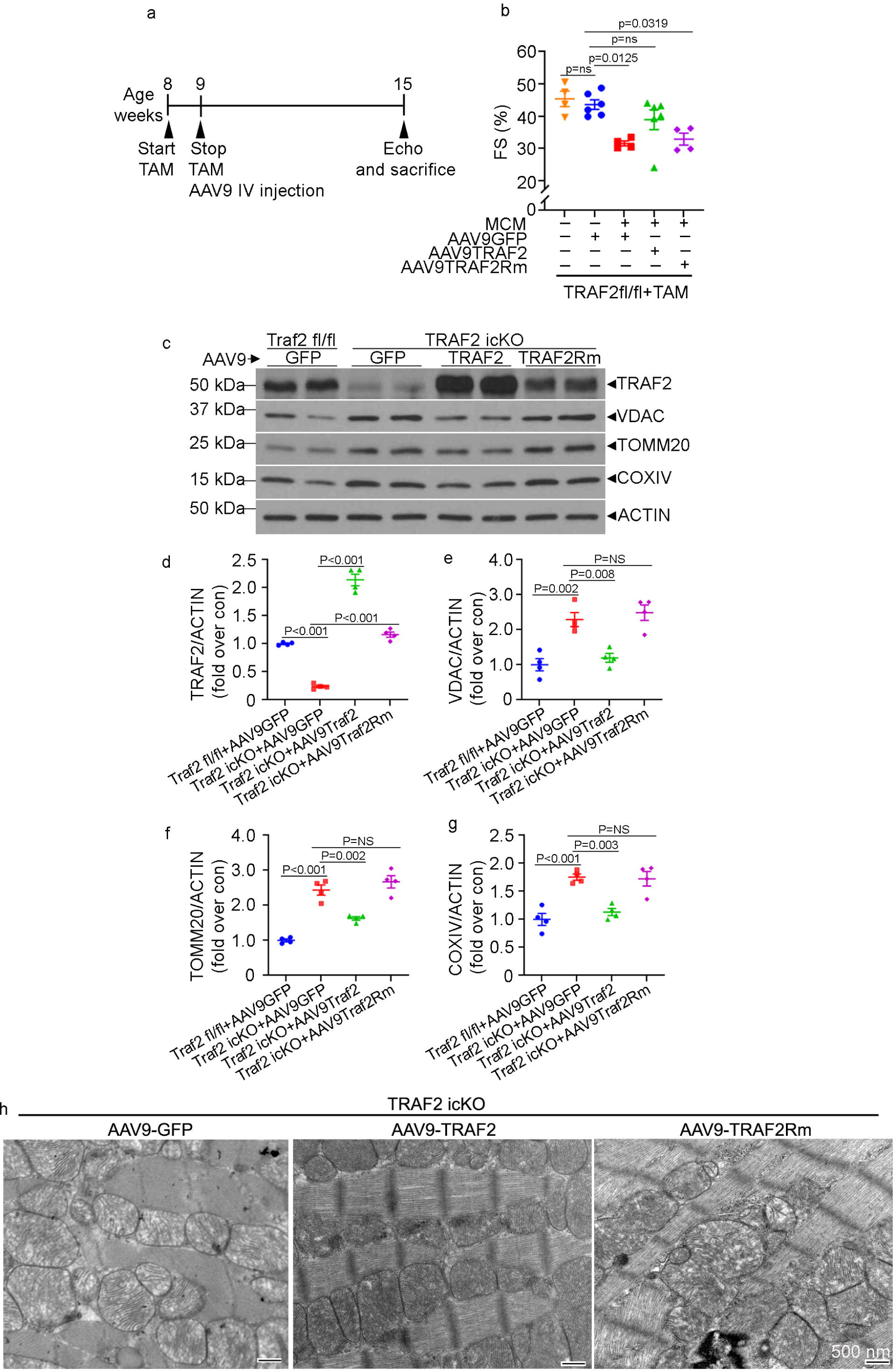
Restoration of TRAF2, but not its E3 ligase deficient mutant, rescues cardiomyopathy and mitochondrial abnormalities with inducible cardiac myocyte TRAF2 ablation. **a.** Schematic depicting experimental strategy for tamoxifen inducible TRAF2 ablation in mice homozygous for floxed *Traf2* alleles carrying *Myh6*-MerCreMer transgene, followed by AAV9-cardiac troponin T promoter-TRAF2 or TRAF2Rm particles at 5.0 × 10^11 viral particles per mouse. **b**. Left ventricular endocardial fractional shortening (%FS) in TRAF2icKO mice or TRAF2 floxed mice transduced with AAV9 particles coding for TRAF2 or TRAF2Rm, driven by a cardiac troponin T promoter as shown in a. P values by post-hoc test after one-way ANOVA. **c-g**. Representative immunoblot (c) with quantitation of TRAF2 expression (d) and abundance of mitochondrial proteins (VDAC, e; TOMM20, f; COXIV, g); all normalized to ACTIN) in cardiac extracts from mice modeled as in a. P values shown are by post-hoc test after one-way ANOVA. **h**. Representative transmission electron microscopy images to evaluate myocardial ultrastructure in mice as in a. Scale bar= 500 nm.

### Parkin expression is not redundant to TRAF2 function; but Parkin-mediated mitophagy can rescue cell death with TRAF2 ablation

In isolated cardiac myocytes, we have previously observed that TRAF2 co-localizes and interacts with Parkin in the setting of ionophore-induced mitochondrial damage.^30^ Moreover, exogenous Parkin was sufficient to rescue the mitochondrial abnormalities observed with TRAF2 ablation.^30^ We examined Parkin levels in TRAF2-icKO mice and did not note them to be significantly regulated (Fig. 3a). To determine whether endogenous Parkin was playing role in the cardiomyopathy observed with TRAF2 ablation, we generated TRAF2-icKO mice in the *PARK2* null (Parkin deficient) background. As shown in Fig. S8a-e, loss of Parkin did not worsen left ventricular dysfunction or dilation or affect histology or fibrosis in TRAF2-icKO mice. These data are consistent with a lack of role for Parkin in transducing basal mitophagy in cardiac myocytes in unstressed young adult mouse hearts.^12, 22^

To determine if the observed mitophagy defect was responsible for increased cell death with TRAF2 ablation, we studied *Traf2* floxed murine embryonic fibroblasts (*Traf2* fl/fl MEFs), wherein TRAF2 was deleted with adenoviral-Cre expression (Fig, S9a, b). Loss of TRAF2 in MEFs resulted in accumulation of damaged mitochondria by ultrastructural examination (Fig. 8a) with impaired mitophagy assessed with mKeima expression (Fig. 8b, c), and cell death (Fig. 8d). Importantly, ablation of TRAF2 did not alter expression of LC3 or p62, two markers of macro-autophagy (Fig. S9b) mirroring the findings in TRAF2-icKO mice (Fig. 3e-g), and confirming a specific role for TRAF2 in mitophagy. Adenoviral-overexpression of Parkin, as we have previously described,^30^ rescued mitophagy to control levels and attenuated cell death with TRAF2 ablation (Fig 8b-d). Taken together, these data indicate that TRAF2 mediates mitophagy independent of Parkin, and the mitophagy defect is the proximate cause of cell death in the absence of TRAF2.

**Figure 8.**
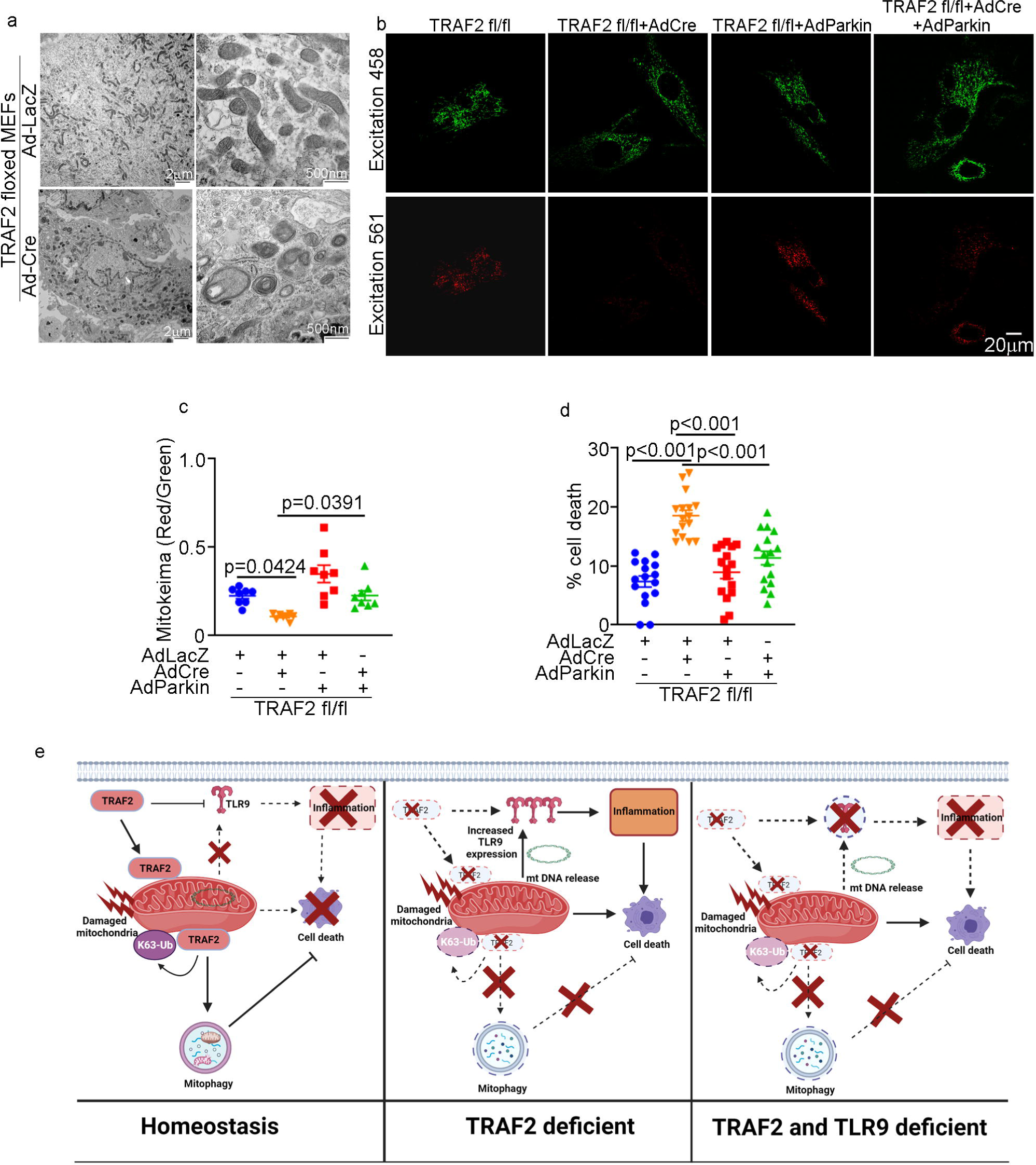
TRAF2 ablation induced cell death, which can be rescued by Parkin-induced mitophagy. **a.** Murine embryonic fibroblasts (MEFs) carrying floxed *Traf2* alleles were modeled for TRAF2 ablation with adenoviral Cre (or LacZ as control, each at MOI=100) treatment for 72 hours as in Fig. S9 and subjected to transmission electron microscopic (TEM) analysis. Representative TEM images demonstrating mirtochondrial abnormalities noted after *Traf2* ablation as compared with control. **b, c**. Representative images (b) and quantitative analyses of mitoKeima emission in the red channel over green as index of mitophagy (c), in *Traf2* null MEFs (modeled as in a) expressing mitoKeima transduced with adenoviral Parkin or LacZ (MOI=100 for 48 hours) as control. Adenoviral particles coding for LacZ were added to equalize the number of viral particles per treatment. P values shown are by post-hoc test after one-way ANOVA. **d**. Cell death in *Traf2* null MEFs (modeled as in a) and transduced with adenoviral Parkin or LacZ (MOI=100 for 72 hours) as control. Adenoviral particles coding for LacZ were added to equalize the number of viral particles per treatment. P values shown are by post-hoc test after one-way ANOVA. **e**. Schematic depicting role of TRAF2-induced mitophagy in preventing inflammation and cell death in cardiac myocyte homeostasis. TRAF2 localizes to the mitochondria and promotes mitophagy via K63-ubiquitination activity in unstressed cardiac myocytes (*left panel*). Ablation of TRAF2 results in mitochondrial DNA leak, and upregulates TLR9 abundance to provoke inflammation via mitochondria DNA sensing. The resultant inflammation and accumulation of damaged mitochondria provoke cardiac myocyte cell death and cardiomyopathy (*middle panel*). Concomitant ablation of TLR9 in the setting of TRAF2 deficiency prevents inflammation and cell death, and rescues left ventricular hypertrophy and function in the short-term; but does not prevent accumulation of damaged mitochondria and cell death inexorably resulting in cardiomyopathy during follow up (*right panel*).

## Discussion

Purported as ancient symbionts, mitochondria act as arbiters of cellular viability and death.^1^ Furthermore, mitochondrial damage triggers inflammatory signaling.^1, 2^ In this work, we have identified the homeostatic role for mitochondrial quality control in cardiac myocytes, and distinguished the effects of inflammation from those due to persistent cell death signaling from damaged mitochondria in the development of cardiomyopathy. Our studies reveal a novel regulatory pathway that links two key elements of the innate immune system, namely TRAF2 and TLR9 to balance mitophagy and suppress inflammatory signaling for maintaining myocardial homeostasis in the unstressed heart (Fig. 8e). Importantly, our data demonstrate that TRAF2, an adaptor protein downstream of receptors that drive innate immunity signaling, localizes to the mitochondria in the unstressed heart (Fig. 1), and loss of TRAF2 impairs homeostatic mitophagy (Fig. 3). Ablation of TRAF2 in adult cardiac myocytes induces cardiac myocyte cell death and cardiomyopathy (Fig. 2). This phenotype is rescued by restoration of the wild type TRAF2 protein, but not a mutant lacking E3 ligase activity (Fig. 7). TRAF2 ablation-induced mitophagy impairment plays a direct role in causing cell death, as rescuing mitophagy with Parkin overexpression prevents cell death downstream of TRAF2 deficiency (Fig. 8). Our data also indicate that TRAF2 acts as a brake to suppress TLR9 signaling, as loss of TRAF2 upregulates mitochondrial DNA sensing via TLR9 to drive local inflammation, which plays a critical mechanistic role early in the observed cardiomyopathy (Fig. 4, 5). Persistence of damaged mitochondria in cardiac myocytes from mice with inducible TRAF2 ablation in a TLR9 deficient background is associated with inexorable progression to cardiomyopathy (Fig. 6), suggesting that facilitating removal of damaged mitochondria is essential and targeting downstream mitochondrial DNA sensing and resultant inflammation only offers temporary reprieve from downstream effects of mitochondrial damage. Overall, these findings indicate that despite a vital role for mitochondria in cellular energy generation, rapid removal of damaged mitochondria via mitophagy is essential to maintaining a pool of normally functioning mitochondria, while preventing activation of cell death pathways and downstream inflammatory signaling. Moreover, the observation that TRAF2 localizes to mitochondria in human myocardium, with increased localization in cardiomyopathic hearts (Fig. 1) offers hope for boosting this pathway to facilitate mitochondrial quality control and stimulate healing.

Cardiac myocytes display robust levels of basal mitophagy in the unstressed heart as assessed by various contemporary techniques,^11, 14, 16, 17, 19, 21^ which protects against activation of apoptotic and/or necroptotic signaling pathways triggered by mitochondrial permeabilization.^1^ Indeed, under conditions of stress when mitochondrial damage is upregulated, such as with pressure overload^9^, myocardial infarction^8, 11^ nutrient excess^10^ or aging^13, 14^, impairment of mitophagy is associated with increased cardiac myocyte death. Mitophagy is also a critical mechanism to prevent sterile myocardial inflammation provoked by sensing of mitochondrial DNA leaked from damaged mitochondria. Indeed, this is particularly relevant as TLR9 signaling has been shown to drive inflammatory signaling in the setting of mitochondrial DNA release, when mitochondria are increasingly damaged, such as with pressure overload-induced stress or with ischemia-reperfusion injury.^2, 4^ Degradation of cytokine mRNAs by Regnase-1 has been demonstrated as another mechanism to prevent sterile inflammation in the myocardium in the setting of pressure overload hypertrophy.^44^ Curiously, while lysosomal DNase2a and Regnase-1 have been shown to be critical for protecting against myocardial inflammation under stress, mice deficient in these genes in cardiac myocytes do not have detectable myocardial inflammation or cardiac structure or function abnormalities in the unstressed state.^2, 44^ These observations strongly support the contention that robust and efficient basal mitophagy removes damaged mitochondria during homeostasis (in the unstressed state), to prevent mitochondrial DNA sensing and sterile inflammation. Our studies point to TRAF2 as the sensor that resides in the mitochondria to effect homeostatic mitophagy upon mitochondrial damage to prevent cardiac myocyte cell death. Indeed, TRAF2-mediated mitophagy prevents mitochondrial DNA release and DNA-sensing to drive inflammation, as demonstrated with attenuation of myocardial CD68+ macrophage abundance to levels seen in unstressed state, in TRAF2-icKO mice lacking TLR9. It is important to note that while TLR9 ablation rescued cardiac myocyte cell death in the short term by preventing mitochondrial DNA-sensing and inflammation, the persistence of damaged mitochondria in the absence of TRAF2-induced mitophagy led to cardiomyopathy over a longer-term follow up. These data underscore the critical importance for removal of damaged mitochondria to maintain cellular homeostasis.

TRAF2, an adaptor protein, has been ascribed critical roles in innate immunity and inflammatory signaling (reviewed in ^45^). Our studies demonstrate a novel role for TRAF2, as the first mediator to be uncovered for transducing basal cardiac myocyte mitophagy in young adult mice; distinct from previously described roles for other pathways in stress-induced mitophagy^8–10, 13–15, 20^ or mitophagy during perinatal cardiac growth and development.^12^ It is important to note the difference between basal and stress-induced mitophagy, where the PINK1-Parkin pathway has been ascribed an important role. Prior studies have noted a critical role for PINK1 (PTEN-induced putative kinase 1), a mitochondrially-targeted serine-threonine kinase, which gets stabilized upon mitochondrial damage and recruits PARKIN^46^, an E3 ubiquitin ligase, that ubiquitinates mitochondrial proteins and targets the damaged mitochondria to autophagosomes.^47^ Our data indicate that TRAF2 localizes to mitochondria is independent of PINK1 in the unstressed state (Fig. S2), and mediates mitophagy independently of endogenous Parkin (Fig. S8). This is consistent with the lack of homeostatic mitophagy defects observed in Parkin and PINK1 deficient myocardium in unstressed young adult mice.^8, 12, 19, 20, 22^ Indeed, TRAF2 was uncovered in a screen for PINK1 and Parkin-independent mitophagy mediators^48^; and inflammatory stimuli were observed to induce TRAF2-mediated mitophagy via ubiquitination of orphan nuclear receptor, Nur77^32^; independently confirming our prior observations that TRAF2 mediates mitophagy in cardiac myocytes.^30^ Consistent with a role for PINK1-mediated mitophagy under stress, we found that PINK1 was essential for recruitment of TRAF2 to mitochondrial fraction upon hypoxia-reoxygenation injury; but did not affect TRAF2’s localization to the mitochondria under unstressed normoxic conditions (Fig. S2).

TRAF2 harbors a coiled-coiled C-terminal interaction domain and a N-terminally located E3-ubiquitin ligase ring domain; and TRAF2 ubiquitinates multiple proteins to activate signaling pathways working.^49^ Our findings also confirm an important role for TRAF2’s E3 ubiquitin ligase signaling function in facilitating mitophagy. Indeed, TRAF2 facilitates K63 ubiquitination,^49^ and K63 poly-ubiquitination has been shown to selectively target proteins for lysosomal degradation.^50^ We propose that the sub-cellular localization of TRAF2 mediates differential signaling through its E3 ligase activity. Our studies demonstrate that a fraction of TRAF2 associates with the mitochondria in unstressed mouse hearts, in human hearts without pre-existing cardiac pathology (Fig. 1) and in isolated cardiac myocytes^30^ (as well as HEK293 cells, see Fig. S1) where its K63-mediated ubiquitination of specific substrates such as Nur77^32^ and hitherto undiscovered targets may facilitate mitophagy. TRAF2-mediated ubiquitination of RIPK1 is also critical for NFκB activation^51^, which is known to occur upon TNF receptor activation and recruitment of cytosolic TRAF2 to the TNR receptor-signaling complex. Interestingly, TRAF2 has been shown to signal via NFκB to suppress transcription of BNIP3^52^, a BH3-only protein that is activated upon stress to induce mitochondrial permeabilization and cardiac myocyte death.^5^ Prior studies have shown that NFκB activation suppresses TLR9 transcription in human cells;^53^ and TRAF2 mediates degradationof TLR signaling pathway proteins through its E3 ligase activity in myeloid cells.^54^ Our data suggest that TRAF2-mediated transcriptional repression via NFκB is likely operative in keeping TLR9 expression in check in cardiac myocytes to prevent mitochondrial DNA sensing while simultaneously facilitating mitophagy to remove damaged mitochondria. Indeed, ablation of TRAF2 in adult cardiac myocyte triggers transcriptional upregulation of TLR9 in cardiac myocytes (Fig. 4d) concomitantly with increased mitochondrial DNA sensing (Fig. 4a) and CD68+ macrophage infiltration as a direct result of TLR9 signaling (Fig. 5a, b).

Notably, our studies demonstrate induction of cellular inflammation with upregulated macrophage abundance in the myocardium of mice with inducible adult-onset TRAF2 ablation, without evidence for systemic inflammation. These findings are in contrast to prior studies with perinatal ablation of TRAF2 in cardiac myocytes which demonstrate induction of systemic inflammation with increased circulating TNF levels (along with other cytokines) that transduces TNF receptor 1 (TNFR1)-mediated activation of cardiac myocyte necroptosis.^34^ Similar observations were made in germline TRAF2 deficient mice^55^ that demonstrate systemic inflammation, markedly elevated TNF-induced cell death and early lethality. On the contrary, ablation of TRAF2 in myeloid cells,^54^ hepatocytes^56^, or B cells^57^ did not induce systemic inflammation, cell death or mortality in unstressed mice. Taken together, these observations argue for the developmental and/or cell-type specific effects of TRAF2 signaling in preventing necroptosis that are not observed in adult cardiac myocytes.

Our findings are consistent with a growing body of literature supporting a cardio-protective role for TRAF2 signaling. Indeed, spurred by initial observations that TNFR receptor signaling conferred cytoprotection in the setting of myocardial infarction,^58^ our lab has demonstrated that TRAF2, a common adaptor downstream of both TNF receptors confers protection against cell death with ischemia-reperfusion injury.^35^ In the current study, we observed upregulation of TRAF2 in human myocardium from patients with ischemic cardiomyopathy with a dramatic increase in its localization to the mitochondrial fraction (Fig. 1b, d). This was observed in murine hearts as well,^30^ and we confirmed that TRAF2 levels increase in purified mitochondria from mouse hearts subjected to ischemia-reperfusion injury (Fig. 1g, h). We surmise that TRAF2 localized to the mitochondrial facilitates stress-induced mitophagy as a likely mechanism for its role in reducing infarct size with complete coronary occlusion^34^; and TRAF2-mediated mitophagy is another potential mechanism for the cytoprotection conferred with TRAF2 overexpression (at modest levels) in murine hearts subjected to ischemia-reperfusion injury.^35, 36^

In summary, the data presented herein identify mitochondrial quality control mediated by TRAF2 as an essential homeostatic phenomenon in the adult myocardium. In doing so, TRAF2 minimizes deleterious intracellular inflammatory signaling, both by preventing the accumulation of depolarized damaged mitochondria, as well as the impact of released mtDNA by suppressing the TLR9 levels to attenuate mtDNA sensing. These data support the premise for augmenting TRAF2 signaling to enhance mitochondrial quality control as a novel therapeutic strategy in cardiomyopathy.

## Methods

### Reagents and samples

We crossed mice homozygous for floxed *Traf*2 alleles (*Traf2* fl/fl)^57^ (which permits Cre-mediated excision of exon 3 and introduces a frame shift resulting in loss of TRAF2 protein; obtained from Dr. Robert Brink, Garvan Institute of Medical Research, Australia) with mice carrying the *Myh6* promoter driven Mer-Cre-Mer (MCM) transgene (generous gift from Jeffery D. Molkentin, Ph.D., Cincinnati Childeren’s Hospital, Cincinnati, OH).^59^ Mice were injected with tamoxifen i.p. or fed with tamoxifen chow (ENVIGO, cat#TD.130857), as described for individual experiments. *Tlr9* null mice were generously provided by Drs. Shizou Akira and Takeshi Saito, Laboratory of Host Defense, Research Institute for Microbial Diseases, Osaka University, Osaka, Japan. *Parkin* null (Strain number 006582), *Pink1* null (stock number 017946) and the autophagic reporter CAG-RFP-EGFP-LC3 (strain number 027139) mice were purchased from Jackson labs. All mice were maintained on a C57BL6 background. Mouse studies were randomized and observers blinded. All animal studies were approved by the Institutional Animal Care and Use Committee (IACUC) at Washington University School of Medicine. Studies on human tissue were performed under an exemption by the IRB at Washington University School of Medicine because only de-identified human samples were used. Human heart tissue obtained from Kenneth Margulies, M.D. at University of Pennsylvania, Philadelphia, PA. Hearts were categorized as either non-failing (no history of heart failure, obtained from non-failing brain-dead donors) or ischemic cardiomyopathy (obtained at time of orthotopic heart transplantation). After in-situ cold cardioplegia, all hearts were placed on wet ice at 4°C in Krebs-Henseleit buffer. Transmural LV samples were obtained from the LV free wall with epicardial fat excluded. The mitoKeima construct was generously provided by Dr. Atsushi Miyawaki, RIKEN brain science institute in Saitama, Japan.

### Echocardiography and myocardial characterization

2D-directed M-mode echocardiography was performed as we have previously described.^60^ Histologic assessment for myocyte registration was performed with hematoxylin-eosin staining, and myocardial fibrosis was assessed with Masson’s trichrome staining, as we have described.^60^ Transmission electron microscopy was performed in cardiac tissue as described.^60^

### Studies with adeno-associated viral vectors

AAV9 particles coding for TRAF2 or its Rm mutant constructs (which we have described previously^30^) driven by the cardiac troponin T promoter for conferring cardiac myocyte selective expression,^42^ were generated by the Hope Center viral vectors core; using AAV backbone constructs generously provided by Dr. Brent French at University of Virginia, Charlottesville, VA.

### Ex-vivo cardiac ischemia-reperfusion injury

Hearts were isolated and perfused as previously described.^36^ After a 20-minute stabilization period, mouse hearts were subjected to no-flow ischemia (time [t] = 0 minutes) for 30 minutes followed by reperfusion for 60 minutes (total t = 90 minutes).

### Adult cardiac myocyte isolation

Adult mouse hearts were subjected to enzymatic digestion to isolate adult cardiac myocytes, as described.^61^ The remaining cellular fraction was isolated as the ‘non-myocyte’ fraction. The cells were subsequently homogenized in RIPA buffer and subjected to SDS-PAGE gel electrophoresis followed by immunoblotting as we have previously described.^30^

### Preparation of murine embryonic fibroblasts

*Traf2* floxed or *Pink1* floxed mice were subjected to timed mating and murine embryonic fibroblasts were prepared from E14 pups following standard protocols.

### Biochemical subcellular fractionation

Mitochondria-enriched and cytosolic fractions were prepared from hearts and cells following the protocols we have previously described.^30^ Biochemical separation of mitochondria rich fraction into mitochondria stripped of other organelles, MAM and ER was performed with Percoll density centrifugation following published protocols.^33^

### In-situ hybridization with RNAscope

All procedures for *Tlr9* transcript detection with fluorescence in-situ hybridization were performed using the RNAScope Multiplex Fluorescent Reagent kit V2 (Cat. No. 323100; Advanced cell Diagnostics) according to the manufacturer’s instructions. RNAScope Probe-Mm-Tlr9, Mouse (Cat. No. 468281; Advanced Cell Diagnostics) and Opal 690 Dye (Cat. No. FP1497001KT; Akoya BioSciences) were used according to the manufacturer’s instructions for fluorescence in-situ hybridization. Desmin staining was performed subsequently to the HRP blocker step of fluorescence in-situ hybridization. Slides were rinsed and covered with 1:50 dilution of goat polyclonal anti-desmin antibody (Cat. No. SC-7559; Santa Cruz) overnight at 4°C. The next morning, slides were rinsed and covered with 1:200 dilution of Alex Fluor 488 donkey anti-goat secondary antibody (Cat. No. A-11055, Invitrogen) for 45 min at room temperature. Slides were rinsed and mounted, stained with DAPI using Antifade Mounting Medium with DAPI (Cat. No. H-1200; Vectashield). Slides were stored at 4°C in the dark until image analysis. Images were captured on Zeiss LSM 700 confocal microscope.

### Immunofluorescence Analysis

We followed the protocol we have previously described.^60^ Paraffin-embedded heart sections (10 µm thick) were subjected to heat-induced epitope retrieval, followed by blocking, and incubated overnight with primary antibody. After serial washes, samples were stained with AlexaFluor594 (Invitrogen) and mounted with fluorescent 4’,6-diamidino-2-phenylindole mounting medium (Vector Labs, H-1200). Confocal imaging was performed on a Zeiss confocal LSM-700 laser scanning confocal microscope using 639 Zeiss Plan-Neofluar 40/1.3 and 63/1.4 oil immersion objectives, and images were acquired using Zen 2010 software. CD68 (Bio-rad, MCA1957), TLR9 (Novus biological, NBP2-24729), PicoGreen from Quant-iT™ PicoGreen™ dsDNA Assay Kit (Thermo Fisher Scientific, P7589). For CD68 staining we used MaxBlockTMAutofluorescence Reducing Reagent Kit (MaxVision Biosciences Inc., cat# MB-M) to block the high autofluorescence background.

### TUNEL staining

Terminal deoxynucleotidyltransferase-mediated dUTP-biotin nick end labeling (TUNEL) was performed on formalin-fixed and paraffin-embedded tissue sections by using the Dead-End fluorometric TUNEL system (Promega, G3250) following the manufacturer’s directions, as previously described.

### Assessment of autophagic flux

We performed autophagic flux analyses by immunofluorescence examination of frozen myocardial tissue from mice with expression of the RFP-GFP-LC3 reporters, as described.^62^ Only puncta associated with cells identified as myocytes based upon visualization of ‘boxcar’ shaped nuclei were counted.

### Assessment of mitophagy

Mice with transduced with AAV9 particles coding for CMV promoter driven mitoKeima, generated by the Hope Center viral vectors core at Washington University, at a dose of 3.5 × 10^11 viral particles per mouse. Mitophagy was assessed with expression of mitoKeima as previously described.^16, 63^ Briefly, live imaging of freshly harvested cardiac slices were performed with Zeiss LSM5 Pascal confocal microscope. The excitation and emission wavelengths was 543 and 580 nm for red fluorescence channel and 458 and 553 nm for green fluorescence channel. Laser power was individualized for each condition to optimize Keima imaging. MEFs were transfected with pcDNA3.1 encoding for mitoKeima construct using GenJet transfection reagent (Sinagen) following manufacturer’s protocols, and mitophagy was assessed as described, using a Nikon A1R confocal system for liver imaging at the Washington University Center for Cellular Imaging (WUCCI). Adenoviruses expressing Cre and rat Parkin were generated as previously described.^30^ The images were quantitated using ImageJ software (NIH).

### Quantitative PCR analyses

Assessment of transcript abundance was performed as described,^60^ using SyberGreen with primers reported in Supplementary Table 1. Assessment of Pink1 transcript expression was performed using TaqMan gene expression assays (Applied Biosystems) for mouse Pink1 (Mm00550827_m1).

### Immunoblotting

Immunoblotting was performed as previously described.^60^ Specific antibodies employed are as follows: LC3 (Novus Biologicals, NB100-2220); p62 (Abcam, ab5416); TRAF2 (Abcam, ab126758), COX IV (Abcam, ab14744); TOMM20 (Sigma, WH0009804M1); VDAC (Cell Signaling, 4661S); PARKIN (Abcam, ab15954); FACL4 (Abcam, ab155282); Calreticulin Antibody #2891; VAPB (Thermo Fisher Scientific, A302-894A); IRE1α (14C10) (Cell Signaling, 3294S) GAPDH (Abcam, ab22555); TLR9 (Novus biological, NBP2-24729); Actin (Sigma, A2066); PINK1 (MRC PPU products and reagents, S774C (DU17570) and S086D (DU34559), and α-Sarcomeric actin (Abcam, ab52219).

### ELISAs

ELISAs were performed on sera collected from mice at terminal bleed with the following kits per manufacturers’ instructions: Mouse IL-1β/IL-1F2 (R&D Systems, MLB00C), mouse TNF-alpha (R&D Systems, MTA00B), mouse IL-6 (R&D Systems, M6000B), and HMGB1 (Chondrex, 6010).

### Immuno-gold detection

For immunolocalization, cells were fixed in 4% paraformaldehyde/0.05% glutaraldehyde (Polysciences Inc., Warrington, PA) in 100mM PIPES/0.5mM MgCl2, pH 7.2 for 1 hr at 4°C. Samples were then embedded in 10% gelatin and infiltrated overnight with 2.3M sucrose/20% polyvinyl pyrrolidone in PIPES/MgCl2 at 4°C. Samples were trimmed, frozen in liquid nitrogen, and sectioned with a Leica Ultracut UCT7 cryo-ultramicrotome (Leica Microsystems Inc., Bannockburn, IL). Ultrathin sections of 50 nm were blocked with 5% FBS/5% NGS for 30 min and subsequently incubated with rabbit anti-GFP (Life Technologies Corporation, Carlsbad, CA.)(1:200) for 1 hr at room temperature. Following washes in block buffer, sections were incubated with 18nm colloidal gold conjugated goat anti-rabbit IgG+IgM (Jackson ImmunoResearch Laboratories, Inc., West Grove, PA)(1:30) for 1 hr. Sections will be stained with 0.3% uranyl acetate/2% methyl cellulose and viewed on a JEOL 1200 EX transmission electron microscope (JEOL USA Inc., Peabody, MA) equipped with an AMT 8 megapixel digital camera and AMT Image Capture Engine V602 software (Advanced Microscopy Techniques, Woburn, MA). All labeling experiments were conducted in parallel with controls omitting the primary antibody.

### Statistical analysis

Statistics were performed in Prism Version 8.0.2 (GraphPad Software Inc.). Statistical significance of differences was calculated via unpaired 2-tailed Student’s t test for 2 group comparisons, or one-way for assessing differences between multiple groups; or 2-way ANOVA to determine mean differences between groups split by 2 independent followed by post-hoc testing to evaluate differences between groups. Graphs containing error bars show means ± SEM with a P value < 0.05 considered statistically significant.

## Acknowledgements

This work was supported by the Hope Center Viral Vectors Core at Washington University School of Medicine. Experiments were performed in part through the use of Washington University Center for Cellular Imaging (WUCCI) supported by Washington University School of Medicine, The Children’s Discovery Institute of Washington University and St. Louis Children’s Hospital (CDI-CORE-2015-505 and CDI-CORE-2019-813) and the Foundation for Barnes-Jewish Hospital (3770 and 4642). The authors are grateful to Jeanne M. Nerbonne, Ph.D., Cardiovascular Division and Center for Cardiovascular Research at Washington University School of Medicine for assistance with adult cardiac myocyte isolation studies. We are also grateful for Wandy Beatty, Ph.D., Department of Molecular Biology at Washington University School of Medicine for assistance with electron microscopy studies. We wish to thank Joan Avery and Katie Kyle at the Center for Cardiovascular Research at Washington University School of Medicine, for their administrative assistance with conducting the study.

