## Supplementary Figures and Table for "TRAF2, an innate immune sensor, reciprocally regulates mitophagy and inflammation to maintain cardiac myocyte homeostasis"

**Supplementary Figure S1: TRAF2 localizes to the mitochondria and mitochondria-associated membranes in unstressed and hypoxia-reoxygenation stressed HEK293 cells.** Representative transmission electron microscopic images in HEK293 cells transduced with GFP-tagged TRAF2 and subjected to immuno-gold detection for GFP under normoxic conditions or after hypoxia-re-oxygenation (24 hours hypoxia followed by 2 hours re-oxygenation) injury. Image from similarly treated un-transfected cells is shown as control. Black arrows point to TRAF2 on mitochondria, white arrows indicate TRAF2 on MAM. Black arrowheads demonstrate TRAF2 localized to a mitochondria within an autophagosome.

**Supplementary Figure S2: PINK1 deficiency does not prevent TRAF2 localization to mitochondria in normoxic culture conditions, but abrogates it with hypoxia-reoxygenation injury. a.** *Pink1* null and wild-type MEFs were subjected to hypoxia-re-oxygenation injury (H/R 24/2 hours) or nomoxia as control followed by biochemical fractionation into mitochondria-enriched (mito, detected by VDAC co-segregation) or cytosolic (cyto, detected with GAPDH expression) fractions. Representative immunoblot depicting detection of TRAF2 in mitochondrial fraction of both groups in normoxic culture conditions, but abrogation of increased TRAF2 abundance in the mitochondrial fraction in WT MEFs by *Pink1* ablation followed hypoxia-reoxygenation injury. **b**. Immunoblot depicting PINK1 in *Pink1* null and wild-type MEFs treated with carbonyl cyanide m-chlorophenyl hydrazone (CCCP, an ionophore that provokes loss of mitochondrial membrane potential and targets mitochondria for autophagic degradation; at 20µM for 24 hours) to stabilize PINK1 followed by immuno-precipitation with anti-PINK1 antibody (MRC PPU products and reagents, S774C (DU17570)) and detection with an alternate PINK1 antibody (S086D (DU34559)) as shown in b. **c**. Detection of *Pink1* transcripts by quantitative PCR analyses in *Pink1* null and wild type MEFs.

**Supplementary Figure S3: Inducible cardiac myocyte TRAF2 ablation in young adult mice results in cardiomyopathy. a**. Representative 2D-directed M mode echocardiographic images from *Traf2* floxed mice carrying the *Myh6*-MerCreMer transgene or with *Traf2* floxed alleles only as control, followed treatment with three doses of tamoxifen i.p. as indicated. Left ventricular end-diastolic diameter (LVEDD) and endocardial fraction shortening (%FS) are reported below the images. N=3/group. * indicates P<0.05 by t-test vs. *Traf2* floxed controls. **b, c**. Representative photograph (b) and Trichrome stained coronal sections (c) from the hearts as in a.

**Supplementary Figure S4: Tamoxifen treatment induces adult onset TRAF2 ablation selectively in cardiac myocytes. a-d.** Representative immunoblots (a, c) demonstrating TRAF2 expression in *Traf2* floxed mice carrying the *Myh6*-MerCreMer transgene or with *Traf2* floxed alleles only (as control) were treated with tamoxifen as indicated. Quantitation is for respective immunoblots is depicted in graphs to the right (b, d). P values are by t-test. **e, f**. Immunoblot (e) and quantitative analyses (f) demonstrating TRAF2 expression in *Traf2* floxed mice carrying the *Myh6*-MerCreMer transgene or with *Traf2* floxed alleles only (as control) 15 days after treatment with tamoxifen (20 mg/kg/d i.p. 5 days/week for 3 weeks), followed by enzymatic digestion to isolate cardiac myocytes from the non-myocyte cellular fraction. Two exposures are shown. P values are by t-test. **g, h**. Representative immunoblot (g) demonstrating TRAF2 expression in *Traf2* floxed mice carrying the *Myh6*-MerCreMer transgene or with *Traf2* floxed alleles only (as control), two weeks after treatment with a single dose of tamoxifen. αSA indicates α-sarcomeric actin (for a, c, g) **i**. Correlation between mean reduction in TRAF2 protein levels and mean % fractional shortening by echocardiographic assessment in TRAF2-icKO mice generated with regimens depicted in a-f. Coefficient of correlation (R) is shown. **j**. Correlation between mean reduction in TRAF2 protein levels and change in left ventricular end-diastolic diameter (LVEDD) from baseline by echocardiographic assessment in TRAF2-icKO mice generated with regimens depicted in a-f. Coefficient of correlation (R) is shown.

**Supplementary Figure S5: Tamoxifen feeding induces adult onset TRAF2 ablation with accumulation of mitochondrial proteins. a.** Immunoblot demonstrating TRAF2 expression in *Traf2* floxed mice carrying the *Myh6*-MerCreMer transgene or with *Traf2* floxed alleles only (as control) two weeks after the mice were fed with tamoxifen chow for the indicated duration in days (D). **b-f**. Immunoblot (b) and quantitative analyses (c-f) demonstrating expression of TRAF2 (b, c), VDAC (b, d), Tomm20 (b, e) and COXIV (b, f) in *TRAF2* floxed mice carrying the *Myh6*-MerCreMer transgene or with *Traf2* floxed alleles only (as control) 2 weeks after the mice were fed with tamoxifen chow for 7 days**.** P values shown are by t-test.

**Supplementary Figure S6: Adult onset inducible TRAF2 ablation does not result in increased circulating markers of cellular necrosis or inflammation. a-d.** Assessment of circulating levels of HMGB-1, IL1β, TNF and IL-6 in sera from mice modeled for inducible cardiac myocyte TRAF2 ablation (TRAF2-icKO) versus TRAF2 floxed mice as controls. Mice treated with a dose of lipopolysaccharide (LPS, 200μg) with sera harvested 90 minutes later were employed as an additional control. '*' indicated P<0.05 by post-hoc test after one-way ANOVA.

**Supplementary Figure S7: TLR9 ablation prevents fibrosis in mice with inducible cardiac myocyte TRAF2 ablation in the short term. a.** Representative immunoblot depicting TRAF2 expression in cardiac extracts from mice with adult-onset inducible TRAF2 ablation (TRAF2-icKO) and TRAF2 floxed controls, without and with concomitant germline TLR9 ablation, 2 weeks after TRAF2 ablation. **b.** Representative immunoblot depicting TLR9 expression in mice with adult-onset inducible TRAF2 ablation (TRAF2-icKO) with and without concomitant TLR9 ablation. **c,d**. Representative images with hematoxylin and eosin staining (c), and trichrome staining (d) to evaluate myocardial structure and fibrosis, respectively, in mice as treated in a. **e,f**. Expression of *myh6* (e) and *myh7* (f) transcripts in mice modeled as in a. No statistically significant differences were detected by one-way ANOVA.

**Supplementary Figure S8: Parkin ablation does not alter cardiac structure or function in mice with inducible cardiac myocyte TRAF2 ablation. a-c.** 2D-directed M-mode echocardiography-derived left ventricular (LV) end-diastolic diameter (LVEDD, a), endocardial fractional shortening (%FS, b), left ventricular mass (LV mass, c) in *Traf2* floxed, *Traf2* floxed bearing MerCreMer transgene (TRAF2-icKO), and *Traf2* floxed bearing MerCreMer transgene in a *Parkin* null background (TRAF2-icKO-Parkin -/-) mice before (PreTAM) and 14 days after tamoxifen treatment for 1 week (postTAM). '*' indicates P<0.05 by post-hoc test after two-way

ANOVA. **d, e**. Representative images with hematoxylin and eosin staining (d), and trichrome staining (e) to evaluate myocardial structure and fibrosis, respectively, in mice as treated in a.

**Supplementary Figure S9. TRAF2 ablation in murine embryonic fibroblasts does not alter macro-autophagy. a.** Murine embryonic fibroblasts (MEFs) carrying floxed *Traf2* alleles were modeled for TRAF2 ablation with adenoviral Cre (or LacZ as control, each at MOI=100) treatment for 24, 48 and 72 hours, with assessment for ablation of *Traf2* gene by PCR analysis. **b**. MEFs treated as above were also subjected to immunoblotting for TRAF2, LC3, SQSTM1 (p62) and GAPDH proteins.

**Supplementary Table 1:** Mouse Primer sequences employed for qPCR analysis of the indicated genes

| Target gene | Forward Primer sequence (5'–3') | Reverse Primer sequence (5'–3') |
|---|---|---|
| *GAPDH* | ACTCCCACTCTTCCACCTTC | TCTTGCTCAGTGTCCTTGC |
| *Acta1* | ACCATCGGCAATGAGCGTTTCC | GCTGTTGTAGGTGGTCTCATGG |
| *Myh7* | GCTGGAAGATGAGTGCTCAGAG | TCCAAACCAGCCATCTCCTCTG |
| *Myh6* | GCTGGAAGATGAGTGCTCAGAG | CCAGCCATCTCCTCTGTTAGGT |
| *Tlr9* | CAAGAACCTGGTGTCACTGC | TGCGATTGTCTGACAAGTCC |
| *Serca2a* | GTGAAGTGCCATCAGTATGACGG | GTGAGAGCAGTCTCGGTAGCTT |
| *Anp* | TACAGTGCGGTGTCCAACACAG | TGCTTCCTCAGTCTGCTCACTC |
| *Bnp* | TCCTAGCCAGTCTCCAGAGCAA | GGTCCTTCAAGAGCTGTCTCTG |

Figure S1

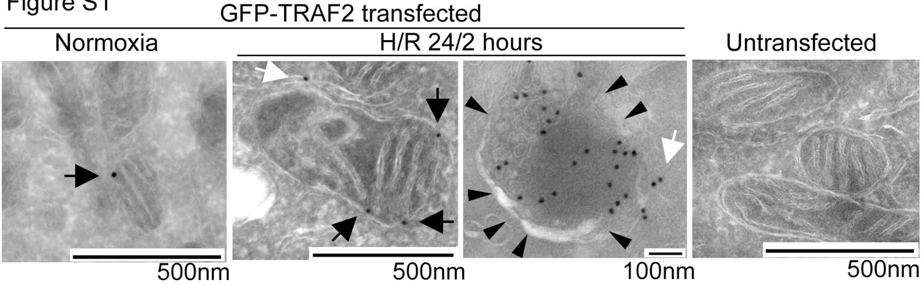
GFP-TRAF2 transfected
Normoxia
H/R 24/2 hours
Untransfected
500nm
500nm
100nm
500nm

Figure S2

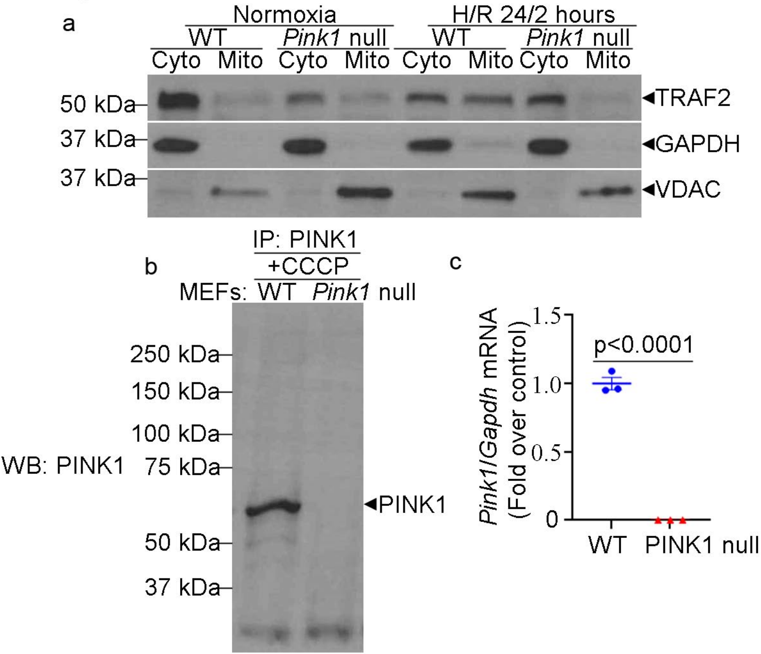

a
Normoxia
H/R 24/2 hours
WT
Pink1 null
WT
Pink1 null
Cyto
Mito
Cyto
Mito
Cyto
Mito
Cyto
Mito
50 kDa
37 kDa
37 kDa
TRAF2
GAPDH
VDAC
b
IP: PINK1
+CCCP
MEFs: WT Pink1 null
WB: PINK1
250 kDa
150 kDa
100 kDa
75 kDa
50 kDa
37 kDa
PINK1
c
Pink1/Gapdh mRNA (Fold over control)
1.5
1.0
0.5
0
p<0.0001
WT
PINK1 null

Figure S3

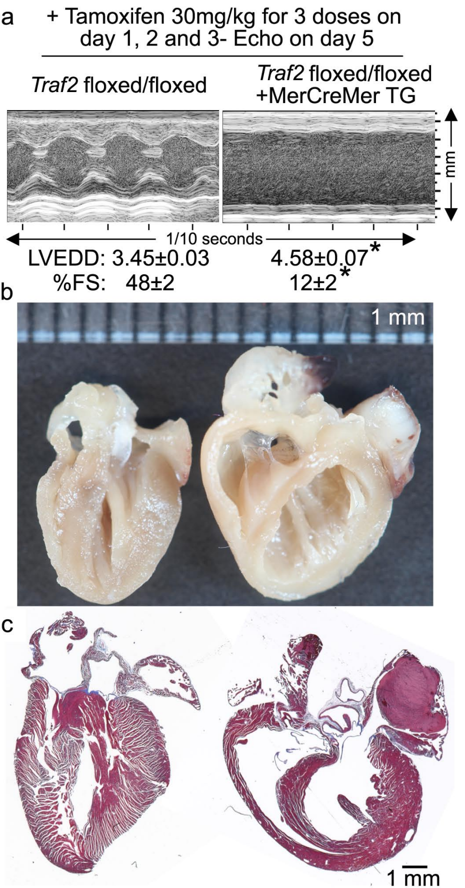
a
+ Tamoxifen 30mg/kg for 3 doses on day 1, 2 and 3- Echo on day 5
Traf2 floxed/floxed
Traf2 floxed/floxed +MerCreMer TG
mm
1/10 seconds
LVEDD: 3.45±0.03
4.58±0.07*
%FS: 48±2
12±2*
b
1 mm
c
1 mm

Figure S4

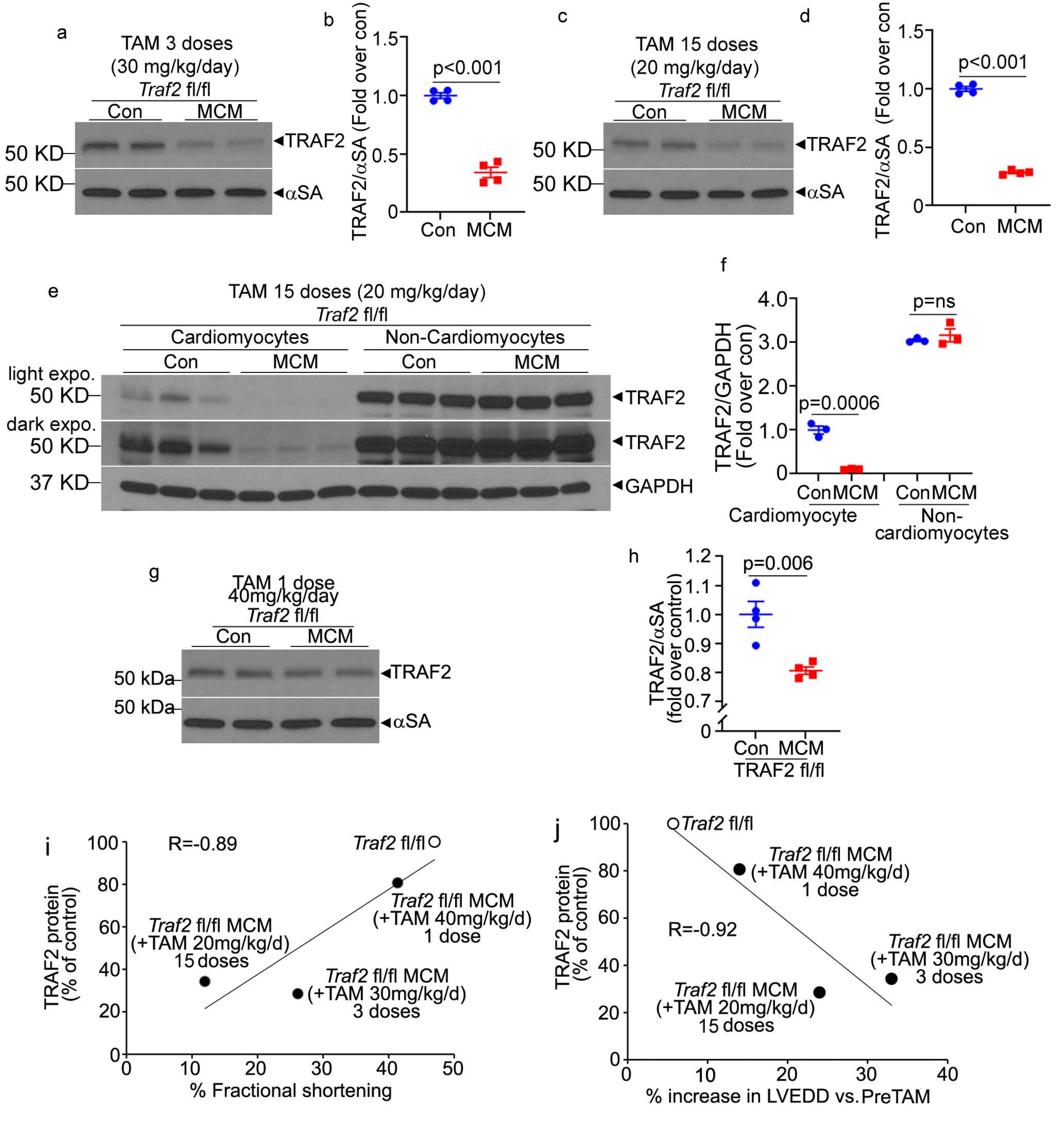

a
TAM 3 doses
(30 mg/kg/day)
Traf2 fl/fl
Con
MCM
50 KD
50 KD
◄TRAF2
◄αSA
b
TRAF2/αSA (Fold over con)
1.5
1.0
0.5
0
p<0.001
Con
MCM
c
TAM 15 doses
(20 mg/kg/day)
Traf2 fl/fl
Con
MCM
50 KD
50 KD
◄TRAF2
◄αSA
d
TRAF2/αSA (Fold over con)
1.5
1.0
0.5
0
p<0.001
Con
MCM
e
TAM 15 doses (20 mg/kg/day)
Traf2 fl/fl
Cardiomyocytes
Non-Cardiomyocytes
Con
MCM
Con
MCM
light expo.
50 KD
dark expo.
50 KD
37 KD
◄TRAF2
◄TRAF2
◄GAPDH
f
TRAF2/GAPDH (Fold over con)
4.0
3.0
2.0
1.0
0
p=ns
p=0.0006
Con
MCM
Con
MCM
Cardiomyocyte
Non-cardiomyocytes
g
TAM 1 dose
40mg/kg/day
Traf2 fl/fl
Con
MCM
50 kDa
50 kDa
◄TRAF2
◄αSA
h
TRAF2/αSA (fold over control)
1.2
1.1
1.0
0.9
0.8
0.7
0
p=0.006
Con
MCM
TRAF2 fl/fl
i
TRAF2 protein (% of control)
100
80
60
40
20
0
R=-0.89
Traf2 fl/fl
Traf2 fl/fl MCM (+TAM 40mg/kg/d) 1 dose
Traf2 fl/fl MCM (+TAM 20mg/kg/d) 15 doses
Traf2 fl/fl MCM (+TAM 30mg/kg/d) 3 doses
0
10
20
30
40
50
% Fractional shortening
j
TRAF2 protein (% of control)
100
80
60
40
20
0
Traf2 fl/fl
Traf2 fl/fl MCM (+TAM 40mg/kg/d) 1 dose
R=-0.92
Traf2 fl/fl MCM (+TAM 30mg/kg/d) 3 doses
Traf2 fl/fl MCM (+TAM 20mg/kg/d) 15 doses
0
10
20
30
40
% increase in LVEDD vs.PreTAM

Figure S5

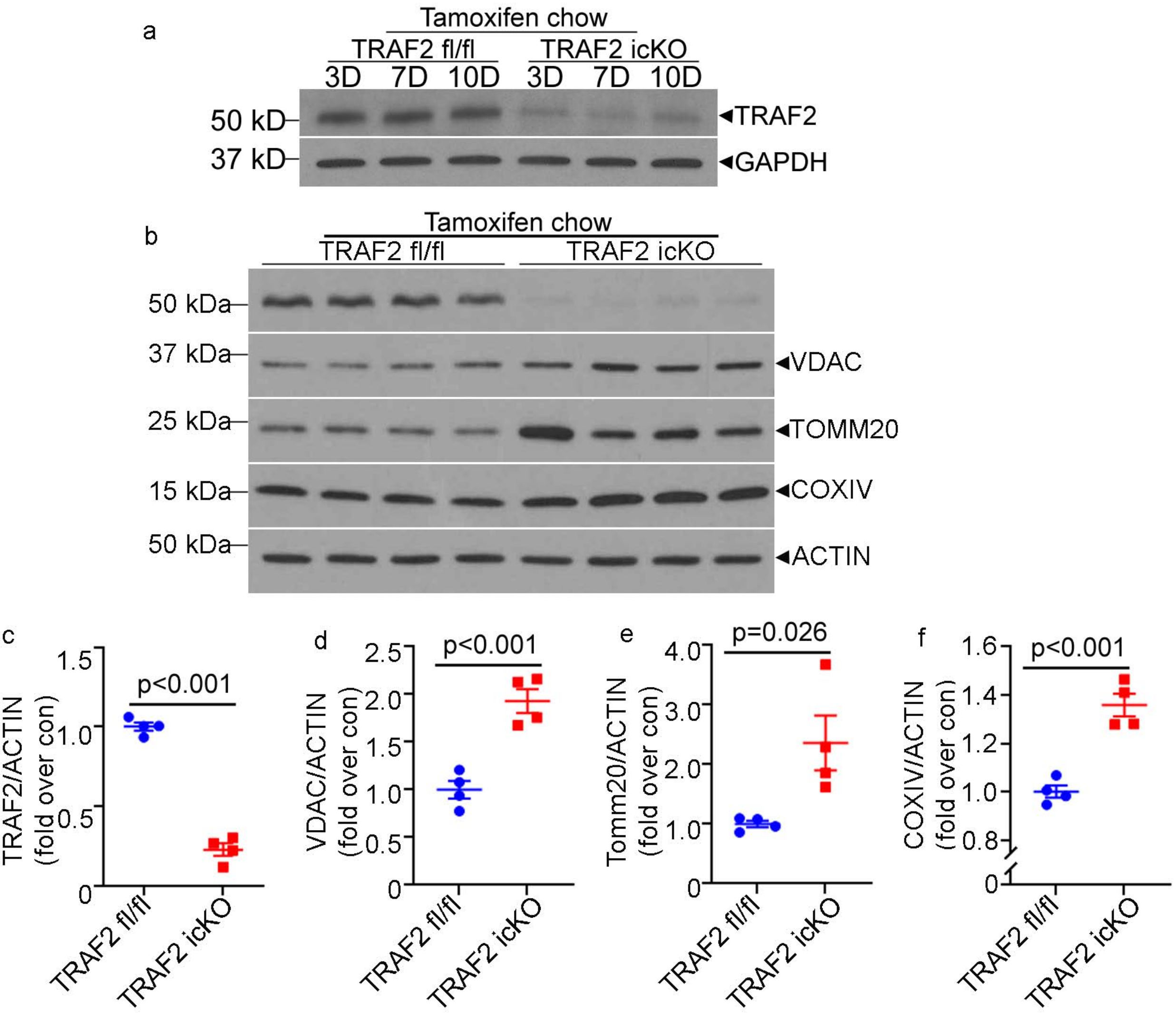
a
Tamoxifen chow
TRAF2 fl/fl
TRAF2 icKO
3D 7D 10D 3D 7D 10D
50 kD
37 kD
TRAF2
GAPDH
b
Tamoxifen chow
TRAF2 fl/fl
TRAF2 icKO
50 kDa
37 kDa
25 kDa
15 kDa
50 kDa
VDAC
TOMM20
COXIV
ACTIN
c
TRAF2/ACTIN (fold over con)
p<0.001
d
VDAC/ACTIN (fold over con)
p<0.001
e
Tomm20/ACTIN (fold over con)
p=0.026
f
COXIV/ACTIN (fold over con)
p<0.001
TRAF2 fl/fl
TRAF2 icKO

### Figure S6

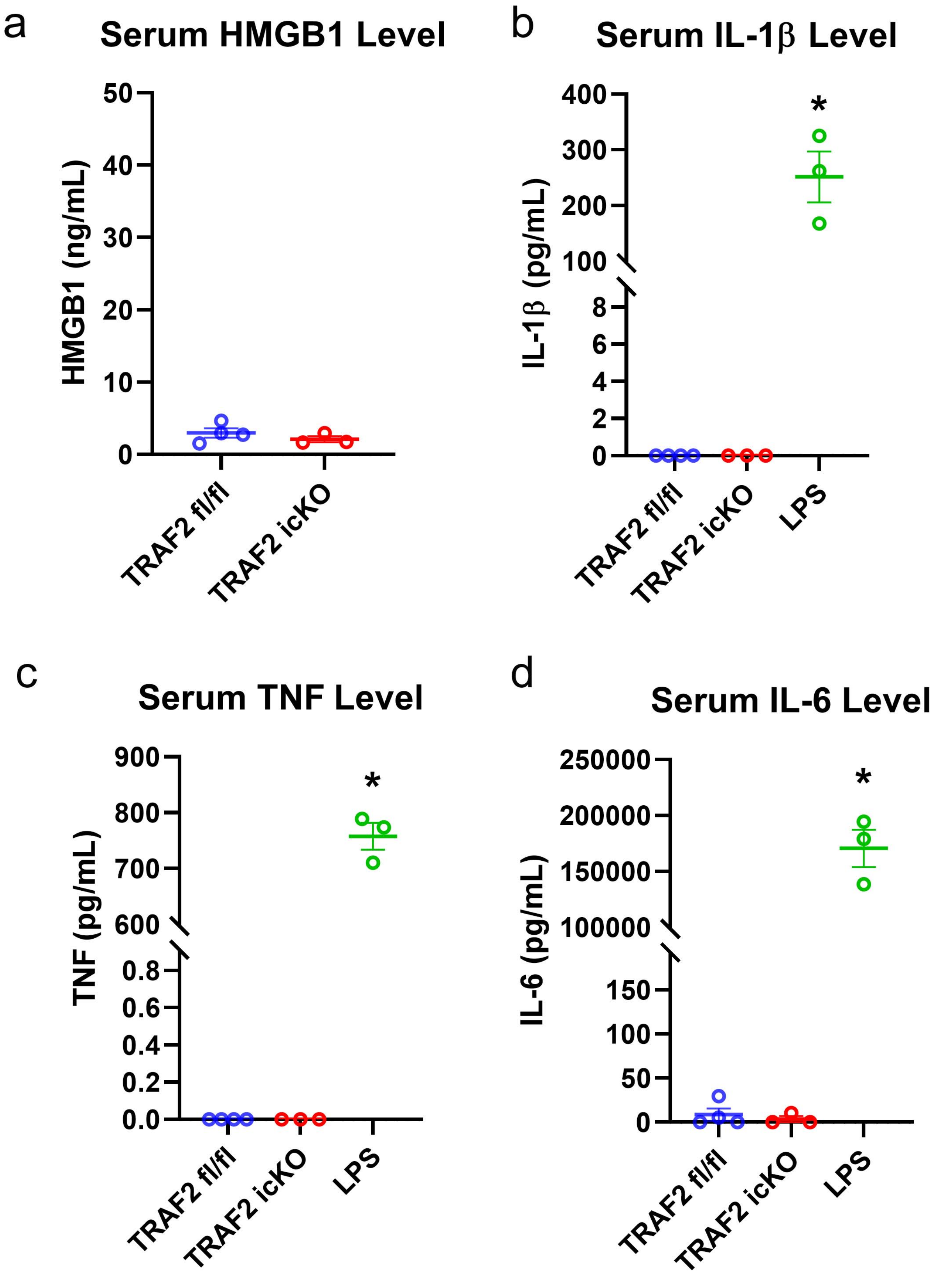
a
Serum HMGB1 Level
HMGB1 (ng/mL)
0
10
20
30
40
50
TRAF2 fl/fl
TRAF2 icKO
b
Serum IL-1β Level
IL-1β (pg/mL)
0
2
4
6
8
100
200
300
400
*
TRAF2 fl/fl
TRAF2 icKO
LPS
c
Serum TNF Level
TNF (pg/mL)
0.0
0.2
0.4
0.6
0.8
600
700
800
900
*
TRAF2 fl/fl
TRAF2 icKO
LPS
d
Serum IL-6 Level
IL-6 (pg/mL)
0
50
100
150
100000
150000
200000
250000
*
TRAF2 fl/fl
TRAF2 icKO
LPS

Figure S7

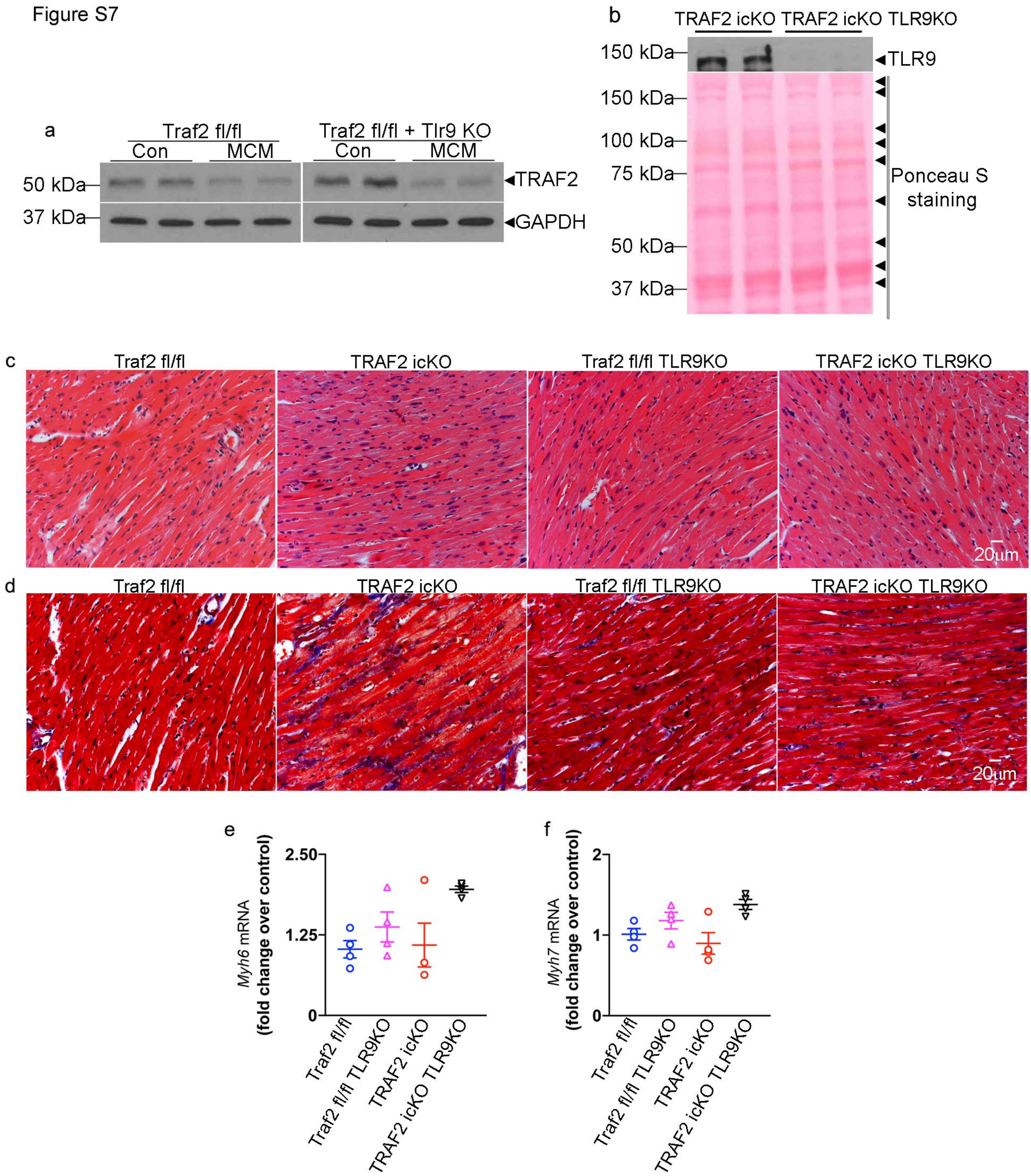
a
Traf2 fl/fl
Traf2 fl/fl + Tlr9 KO
Con
MCM
Con
MCM
50 kDa
37 kDa
TRAF2
GAPDH
b
TRAF2 icKO
TRAF2 icKO TLR9KO
150 kDa
TLR9
150 kDa
100 kDa
75 kDa
50 kDa
37 kDa
Ponceau S staining
c
Traf2 fl/fl
TRAF2 icKO
Traf2 fl/fl TLR9KO
TRAF2 icKO TLR9KO
20μm
d
Traf2 fl/fl
TRAF2 icKO
Traf2 fl/fl TLR9KO
TRAF2 icKO TLR9KO
20μm
e
Myh6 mRNA (fold change over control)
2.50
1.25
0
Traf2 fl/fl
Traf2 fl/fl TLR9KO
TRAF2 icKO
TRAF2 icKO TLR9KO
f
Myh7 mRNA (fold change over control)
2
1
0
Traf2 fl/fl
Traf2 fl/fl TLR9KO
TRAF2 icKO
TRAF2 icKO TLR9KO

Figure S8

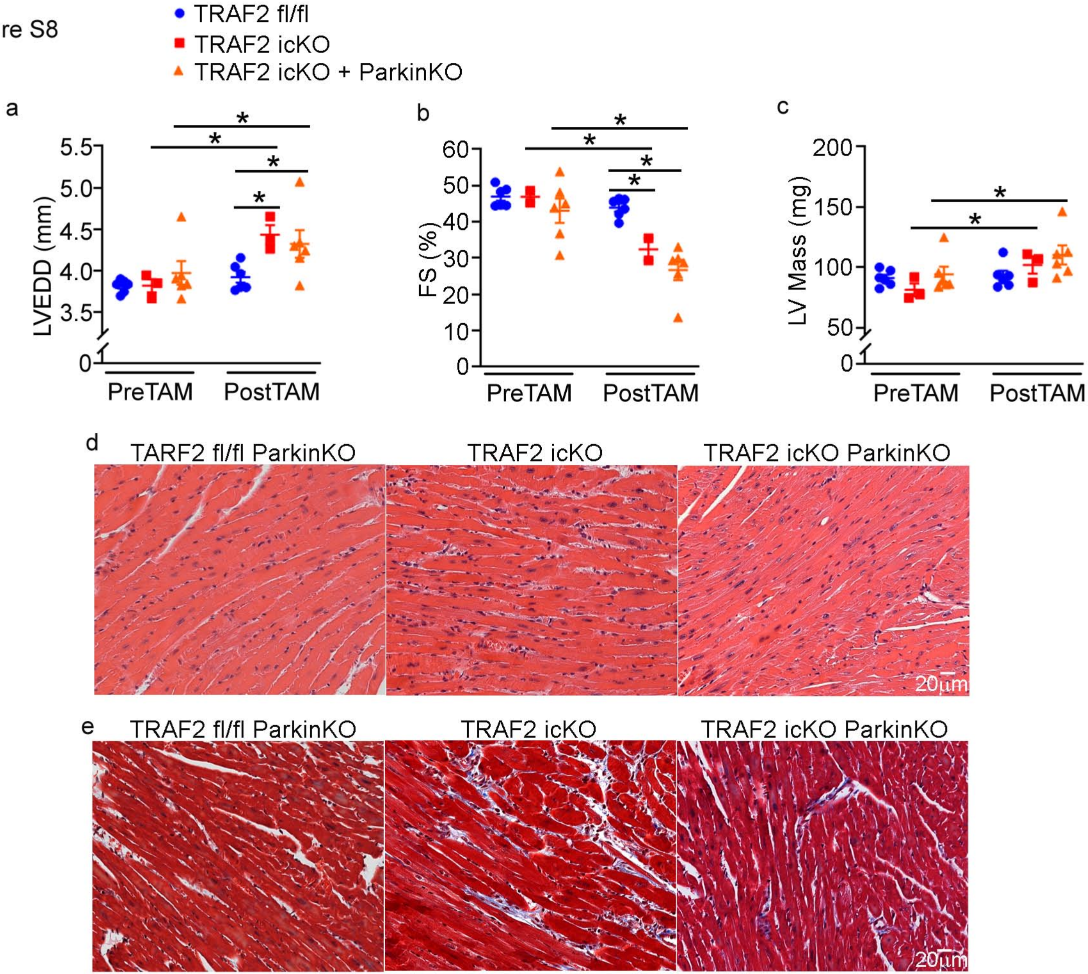
TRAF2 fl/fl
TRAF2 icKO
TRAF2 icKO + ParkinKO
a
LVEDD (mm)
5.5
5.0
4.5
4.0
3.5
0
PreTAM
PostTAM
b
FS (%)
60
50
40
30
20
10
0
PreTAM
PostTAM
c
LV Mass (mg)
200
150
100
50
0
PreTAM
PostTAM
d
TARF2 fl/fl ParkinKO
TRAF2 icKO
TRAF2 icKO ParkinKO
20μm
e
TRAF2 fl/fl ParkinKO
TRAF2 icKO
TRAF2 icKO ParkinKO
20μm

Figure S9

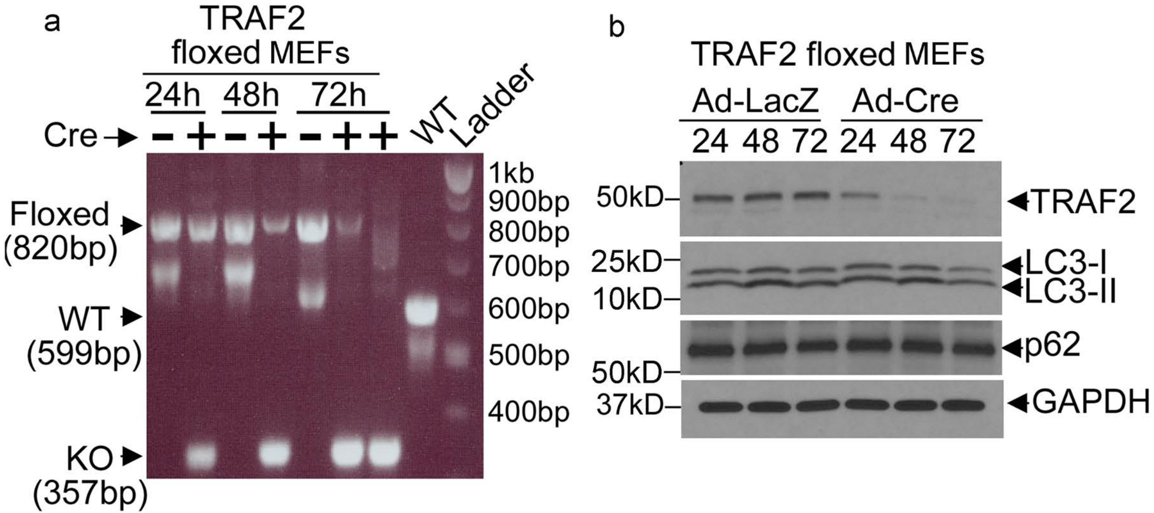
a
TRAF2
floxed MEFs
24h
48h
72h
Cre
-
+
-
+
-
+
+
WT
Ladder
Floxed
(820bp)
WT
(599bp)
KO
(357bp)
1kb
900bp
800bp
700bp
600bp
500bp
400bp
b
TRAF2 floxed MEFs
Ad-LacZ
Ad-Cre
24
48
72
24
48
72
50kD
25kD
10kD
50kD
37kD
TRAF2
LC3-I
LC3-II
p62
GAPDH
